# Selective Laser Sintering of a Photosensitive Drug: Impact of Processing and Formulation Parameters on Degradation, Solid-State, and Quality of 3D Printed Dosage Forms

**DOI:** 10.1101/2021.04.09.439089

**Authors:** Rishi Thakkar, Daniel A. Davis, Robert O. Williams, Mohammed Maniruzzaman

## Abstract

This research study utilized a light-sensitive drug, nifedipine (NFD), to understand the impact of processing parameters, and formulation composition on drug degradation, crystallinity, and quality attributes (dimensions, hardness, disintegration time) of selective laser sintering (SLS) based 3D printed dosage forms. Selective laser sintering (SLS), in most cases, uses an ultraviolet laser source, and drugs tend to absorb radiation at varying intensities around this wavelength (455 nm). This phenomenon may lead to chemical degradation, and solid-state transformation, which was assessed for nifedipine in formulations with varying amounts of vinyl pyrrolidone-vinyl acetate copolymer (Kollidon® VA 64) and potassium aluminum silicate-based pearlescent pigment (Candurin®), processed under different SLS conditions in the presented work. After preliminary screening Candurin®, surface temperature (ST), and laser speed (LS) were identified as the significant independent variables. Further, using the identified independent variables a 17-run, randomized, Box-Behnken design was developed to understand the correlation trends and quantify the impact on degradation (%), crystallinity, quality attributes (dimensions, hardness, disintegration time) employing qualitative and quantitative analytical tools. The design of experiments (DoE) and statistical analysis observed that LS and Candurin® (%wt) had a strong negative correlation on drug degradation, hardness, and weight, whereas ST had a strong positive correlation with, drug degradation, amorphous conversion, and hardness of the 3D printed dosage form. From this study, it can be concluded that formulation and processing parameters have a critical impact on stability and performance; hence these parameters should be evaluated and optimized before exposing light-sensitive drugs to the SLS processes.

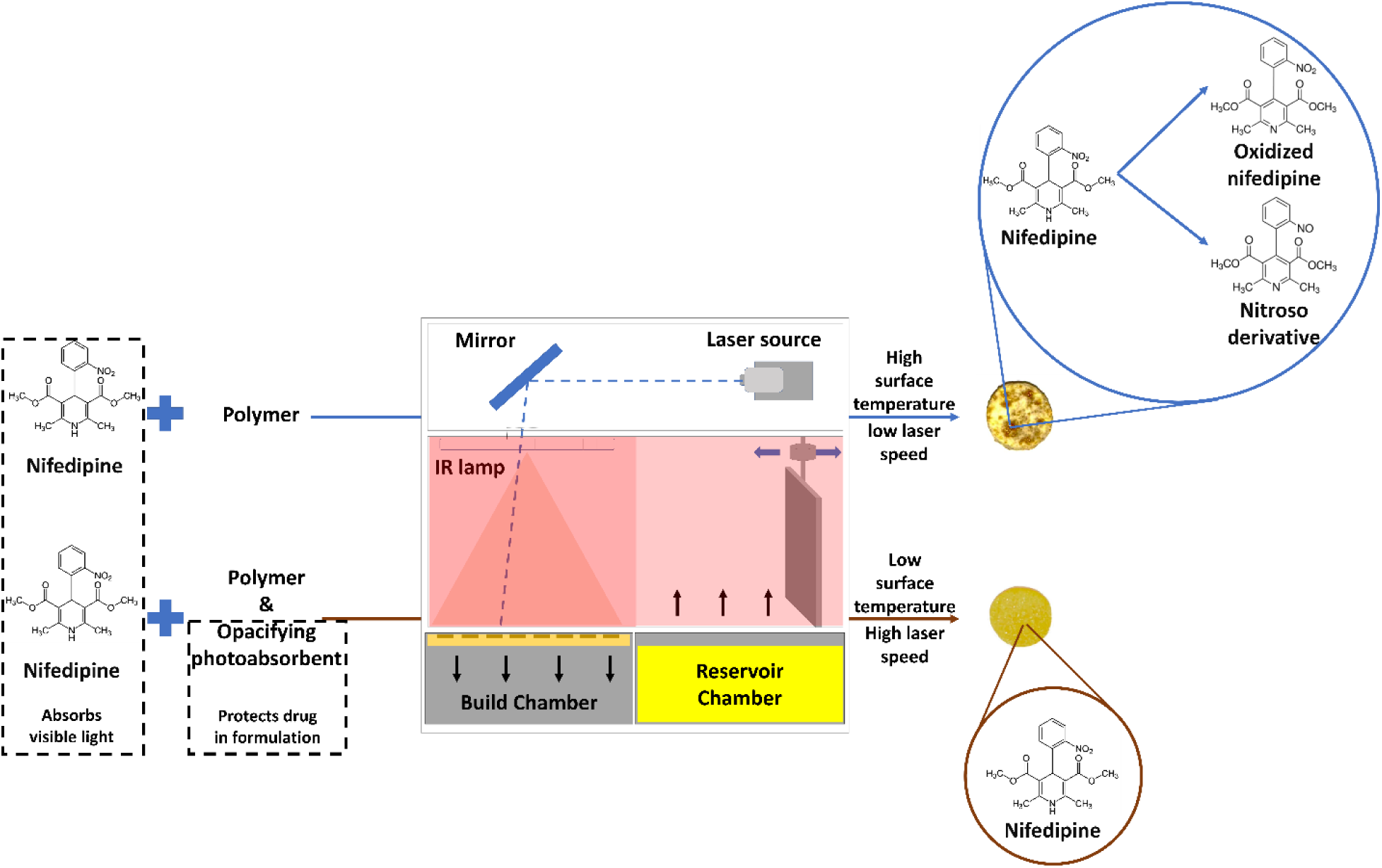

## 1 Introduction

Powder bed fusion (PBF) based 3D printing utilizes either a laser or an electron beam to fuse the feedstock layer by layer into a part^1^. This fusion is attributed to the melting or sintering of the components present in the feedstock, where melting includes heating, liquid conversion, fusion, and solidification, and sintering includes heating and fusion without liquid conversion^2–4^. Thereby, a 3D printing process utilizing a laser source meant to sinter is known as selective laser sintering (SLS)^5^. An SLS process like other powder-bed-based processes utilizes a hopper or a reservoir containing the feedstock (build material), which is dispensed onto the build platform utilizing a roller or a blade in a controlled fashion to maintain the required layer thickness set^6, 7^. The fresh feedstock is spread over the build surface and the high-power laser either sinters or systematically melts the material (x, y, z) described by the underlying g-code^8^. Traditional PBF processes are carried out for thermoplastic polymers and metallic or ceramic composites to build machine parts and tools, where the processing temperature is maintained between the melting temperature (T_m_) and half of the melting temperature (T_m_/2) in a sintering process, and at or above melting temperature in a melting process^9^. Sintering can be the result of multiple chemical and physical processes; diffusion is the most important phenomenon in the process^4,10^. Phase transition after absorbing the laser leads to diffusion and neck formation between the adjacent particles^4,11^. This is the main component of sintering that leads to the lowering of free energy as particles fuse^1, 4, 10^. Moreover, efficient sintering is also dependent on other processing parameters apart from the laser source used^12, 13^. Crucial processing parameters include laser speed, surface temperature, and chamber temperature, whereas print parameters include the layer height, number of perimeters, perimeter offset, hatching offset, and hatching spacing^13, 14^. Moreover, material properties such as flowability impact the print quality and reproducibility for such powder-bed-based platforms, considering the precise layering and mass transfer required in the process^7, 15, 16^.

Owing to the versatility of additive manufacturing processes for rapid prototyping (RP)/ rapid tooling (RT), dosage form designing, and the porosity induced by such powder-bed processes, SLS has found application in the manufacturing of pharmaceutical dosage forms^3^. Previous studies depicting the use of SLS in dosage form development have revolved around immediate release dosage forms^17^, orally disintegrating tablets^18^, modified released dosage forms, and multi-drug releasing systems^19^. Apart from modifying the performance using formulation approaches, researchers have also demonstrated a change in the performance of the dosage forms by merely adjusting the processing parameter^20^. Moreover, this heat-based fusion technique has also been used for the one-step manufacturing of amorphous solid dispersions (ASD) of thermolabile species^14^. Previous research has demonstrated that at lower drug loads (<20%), physical mixtures of the drug in polymeric matrices can be processed at or below the T_g_ of the polymer, to form an ASD, where parameters including hatch spacing, print speed, and surface temperature were found to play a critical role^14^. Processing thermolabile drugs using an SLS process is possible as, unlike selective laser melting, SLS operates at lower temperatures and uses a thermally conductive species or a photo absorbing species for sintering particles together^4^. So far conductive metal particulates such as carbonyl iron or aluminum and photo-absorbing pharmaceutical-grade colorants such as Candurin® have been used to induce sintering in pharmaceutical blends as other pharmaceutical excipients such as polymers might not absorb the radiation or conduct it efficiently^11, 17, 21^.

Bearing in mind the mechanism of SLS 3D printing where a concentrated laser beam is used to sinter material together, it is important to determine the impact of this laser beam and its exposure on the chemistry and the solid-state of the drug^22^. Although most pharmaceutical drugs absorb UV radiation at high intensities below 380nm, there are some which absorb visible radiations over 380 nm and are most likely photolabile^23^. If the chemistry of the molecule is sensitive to light and contains functional groups which may undergo photolysis, they might be susceptible to degradation when exposed to an SLS process although such a study has not been conducted so far. Moreover, most photo-absorbing species are metallic and have high melting points, thereby they do not show any solid-state transformation although the tendency of pharmaceutical small molecules to absorb radiation from the laser source might also impact the solid-state of the molecule. Previous studies exhibiting partial or complete amorphous conversion of the drug have attributed this to the conductive effect of the added sintering agent, and the glass transition of the polymer, although the impact on the solid-state in the absence of a sintering agent or presence of a small molecule which is capable of absorbing the radiation is yet to be inspected^3^.

Nifedipine is a light-sensitive, potent calcium channel blocker used for the treatment of hypertension and angina^23^. Previous studies have confirmed its photolytic degradation through an intra-molecular phenomenon into nitro- and nitroso-pyridine analogs when exposed to ultraviolet and visible light up to 450 nm^24^. Even though the drug depicts two absorption maxima one at 230 nm and the second one at 333 nm nowhere close to the wavelength of the laser, it may absorb the visible radiation subjected to the material during the SLS printing process to some degree, which may lead to the degradation of the drug^25^. This photosensitivity of nifedipine was the primary reason to use it as a model drug for this study. The secondary reason was the biopharmaceutical classification of the drug i.e., BCS class II. Nifedipine has poor aqueous solubility (4-5 µg/mL, pH 7.0), and thereby poor bioavailability which ranges from 40-77% for the immediate release dosage forms *in vivo*^26^. Considering the possibility of solid-state transformation (crystalline to amorphous) of the drug due to its potential sensitivity to the laser source, and the solubility advantage of amorphous systems in comparison to crystalline systems, this study also aimed to demonstrate the SLS-3D printing mediated performance enhancement of a photo-sensitive drug.

Formulations containing the drug (nifedipine (5%-10% w/w)), polymeric carrier (vinyl pyrrolidone-vinyl acetate copolymer (Kollidon® VA 64) (60%-90% w/w)) and sintering agent (potassium aluminum silicate-based pearlescent pigment (Candurin®) (0-30% w/w)) were used for this study. (a) To demonstrate the impact of laser on the degradation and the solid-state of the drug, a physical mixture of the drug and polymer was sintered below the T_g_ of the polymer without the Candurin®. (b) Further, screening studies at 10% w/w drug load with varying amounts of Candurin® and VA64 were conducted to determine the critical formulation and processing parameters. (c) After the preliminary screening, a design of experiments (DoE) study was conducted to determine the impact of the isolated parameters on the degradation and solid-state of the drug at 5% w/w drug loading. High-performance liquid chromatography equipped with a mass spectrophotometric detector (HPLC-MS) and powder X-ray diffraction (pXRD) were used to identify the degradation products and the solid-state characteristics, respectively. Moreover, high-performance liquid chromatography with a UV-Visible detector (HPLC-UV/Vis) was used to quantify the degradation product for the design of experiments (DoE). The quality attributes of the dosage forms, including the dimensions, weight variation, disintegration time, and hardness were also evaluated and used to select an optimized dosage form for *in vitro* performance testing.

## 2 Experimental Section

### 2.1 Materials

The drug, nifedipine, was purchased from Nexconn Pharmtech Ltd. (Shenzhen, China). The polymer, Kollidon®VA64 (average molecular weight 65,000 g/mol), was donated by BASF Corporation (Florham Park, NJ). The electromagnetic energy-absorbing excipient, Candurin®, was purchased from EMD Performance Materials (Philadelphia, PA). Sodium phosphate monobasic, sodium hydroxide, and sodium chloride were purchased from Fisher Scientific (Pittsburgh, PA) for buffer preparation. For dissolution, the bio-relevant fasted state simulated intestinal fluid (FaSSIF) powder was purchased from Biorelevant.com LTD (Surrey, United Kingdom). The selective laser sintering 3-Dimensional desktop printer kit was purchased and self-assembled from Sintratec AG (Brugg, Switzerland). HPLC grade methanol and acetonitrile were purchased from Fisher Scientific; all other chemicals and reagents used were ACS grade or higher.

### 2.2 Preliminary screening and design of experiments

The first step of this study was to determine whether nifedipine (NFD) absorbed visible radiation at a wavelength (λ) of 455 nm, which corresponded to the wavelength of the visible laser-equipped in the selective laser sintering kit (Sintratec kit, Sintratec, Switzerland). For this purpose, a UV-Visible spectrophotometer was used and the absorption spectra of NFD were evaluated. Further, to understand the critical processing and formulation parameters, a screening study was conducted, and the optimal parameters were determined. These optimal parameters and their impact were further evaluated using a design of experiments (DoE) approach. This section of the methods discusses the preliminary screening experiments and the DoE used for this study.

#### 2.2.1 UV-Visible screening studies

Different NFD concentrations were prepared (20, 40, 80, 160 µg/mL) using methanol as a solvent, and their respective absorbance spectrum was collected using a UV-Visible spectrophotometer (Agilent Cary 8454 UV-Vis Diode Array System, Agilent Technologies, Santa Clara, CA). Considering the concentration-based limitations of liquid state quantitative analysis as per Lambert-beer’s law, qualitative investigation of NFD was conducted using a UV-vis reflectance probe with a 316L Stainless Steel/Nickel alloy tip and sapphire window, which was developed to analyze the absorbance of solid samples. The prime objective of this study was to observe the absorbance behavior of NFD around 455 nm. For this experiment, the polymer’s absorbance was not evaluated, as previous studies have demonstrated that it does not absorb visible radiation.

#### 2.2.2 Critical parameters determination

The first step of this study was to determine whether NFD was experimentally absorbing the laser from the source and the laser’s impact on the drug molecule. A physical mixture with Kollidon® VA64 and NFD was subjected to a selective laser sintering process at three different conditions, as depicted in **Table 1**. Post-processing, the printed tablets (printlet) were physically assessed for signs of sintering and were subject to qualitative determination of degradation of the drug post-processing using high-performance liquid chromatography equipped with a mass spectrophotometer (HPLC-MS). Once the laser’s impact on the drug was assessed, a preliminary screening study was conducted to determine the critical formulation and processing parameters. Screening studies are essential to determine and set the range of parameters under investigation for optimization studies. Formulations with a 10% w/w NFD drug loading in different concentrations of Candurin® and Kollidon® VA64 were subjected to SLS 3D printing processes with varying processing parameters (surface temperature, chamber temperature, and print speed). It is important to note that the influence of print parameters (layer height, number of perimeters, perimeter offset, hatching offset, and hatching spacing) was not evaluated as a part of this study and hence were kept constant for all formulations and processing conditions. The formulations and the processing parameters for the screening studies are enlisted in **Table 1**. For the screening studies, the impact of the parameters on the drug’s degradation, amorphous conversion, and, most of all, printability of the drug was assessed. Based on the printability of the printlet the range of the parameters was established for further optimization studies using DoE.

**Table 1.**
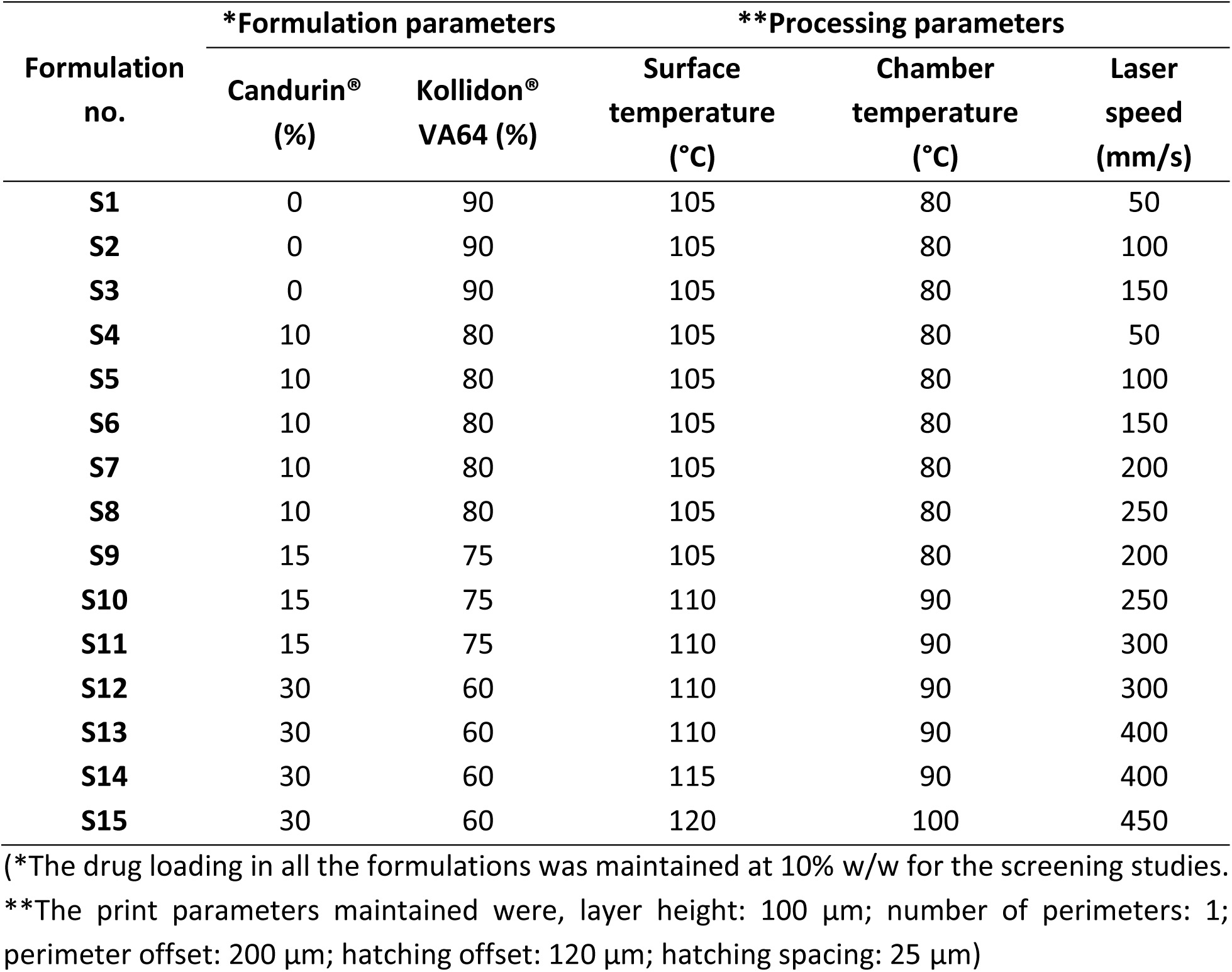
Formulation composition and printing parameters for screening studies.

#### 2.2.3 Optimization studies

After determining the range for the formulation and processing parameters, a response surface DoE study with a 17-run Box-Behnken design was developed using Design-Expert software (Version 10.0.8.0, Stat-Ease, Inc., Minneapolis, MN, USA) to understand the impact of these parameters on the quality attributes (dimensions, weight variation, hardness, disintegration time, density), stability (% degradation), and crystallinity of the printlet and NFD, respectively. For the design, Candurin®(%), surface temperature (°C), and laser speed (mm/s) were considered as the independent variables, whereas crystallinity, degradation (%), hardness, average weight (mg), density (mg/cm^3^), and disintegration time were identified as the dependent variables. The batch to batch variation and reproducibility of the design AM process were assessed by introducing central points in the design, which were repeated five times. The detailed designs and demonstration of variables are shown in **Fig. 1** and **Table 2**.

**Figure 1.**
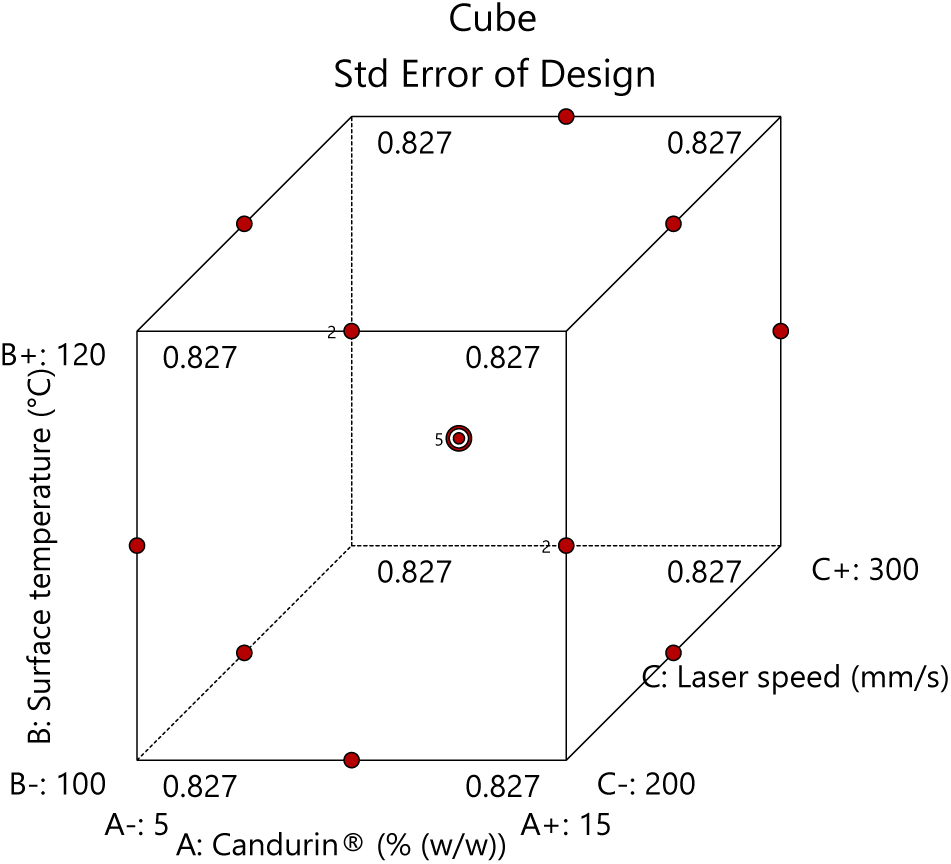
Design points in the Box-Behnken design.

**Table 2.**
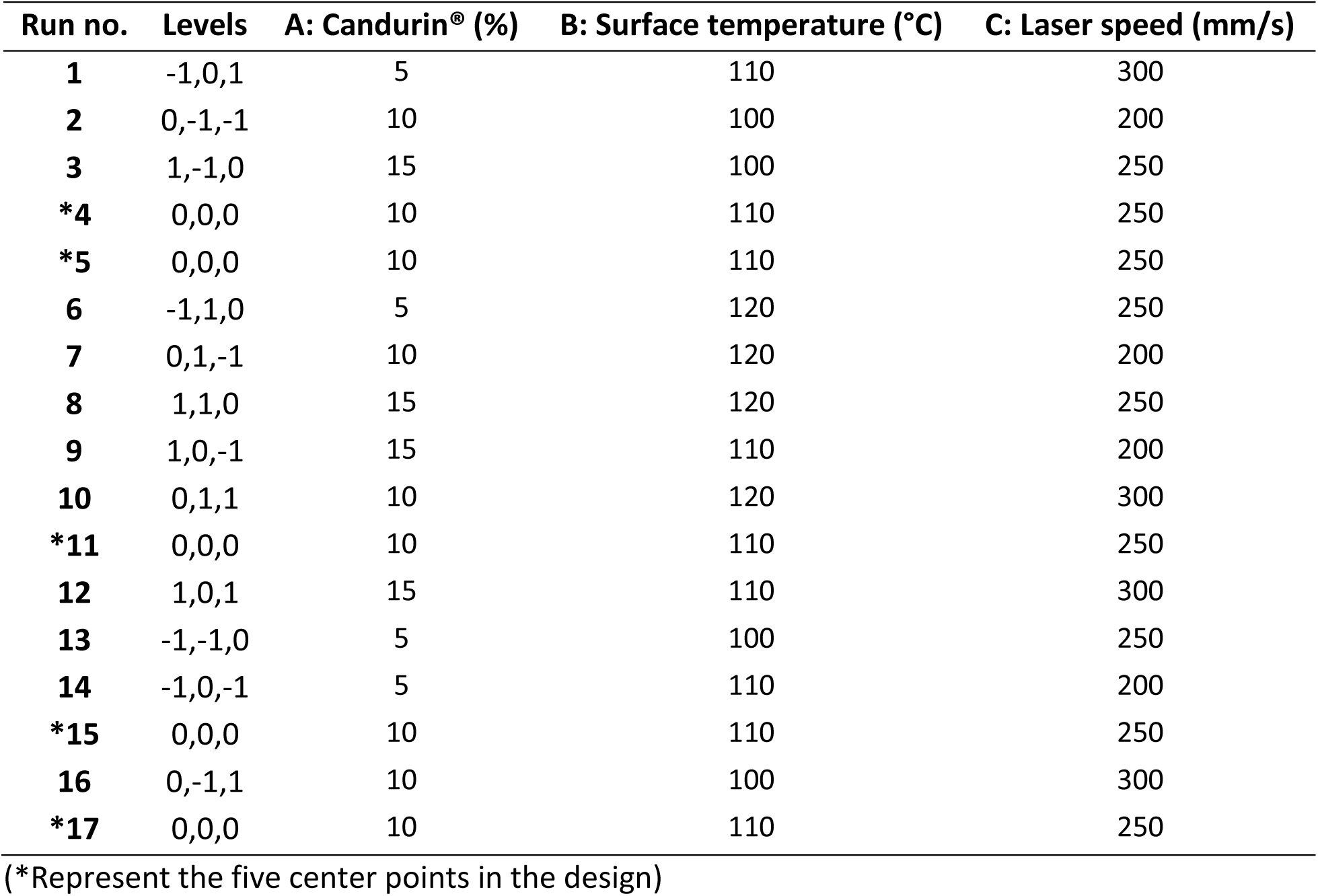
Box-Behnken design for the optimization studies.

The drug loading for the optimization studies was set to 5%, although the ratio (wt%) of NFD to Candurin® in the formulation was maintained as per the screening studies.

### 2.3 Feedstock preparation

Powder-bed-based printers have certain limitations, including but are not limited to the large quantities of feedstock required for the printing process since the powder bed supports the structure being printed. From previous studies without modifying the print bed, typically 150-200 g of feedstock is required based on the dimensions of the printlet, although the un-sintered powder can be recycled. The powder volume can be estimated based on the layer height of the print and the number of layers required to print the part. The second limitation is the absence of mixing of the powder blend during the process. Considering pharmaceutical feedstocks are physical mixtures of multiple components with different densities and bulk properties blended in different ratios, the flow properties of this feedstock play a critical role in the quality attributes of the printlet. Physical mixtures containing NFD, Kollidon® VA 64, and Candurin® were prepared using the geometric dilution technique based on the compositions specified in **Table 1** and **Table 2** for the screening and optimization studies, respectively.

Further, the prepared feedstocks were then passed through the 12-inch diameter, no. 170 sieve (90 µm pore size) to break down any agglomerates present. It should be noted that the sieve pore size should not be more than 100 µm as in that case agglomerates greater than the 100 µm may exist in the feedstock and might be discarded during the printing process instead of being deposited onto the build surface since the layer thickness set for the process is 100 µm. The physical blends were analyzed for drug purity before the process to assess the impact of the process on the degradation of the drug in the blend.

### 2.4 Powder-bed fusion processing (SLS 3D printing)

The feedstock for each screening formulation or optimization run was exposed to PBF based SLS 3D printing process. This powder batch post sieving was added to the feed region of the benchtop LS 3D printer (Sintratec kit, Sintratec, Switzerland). This SLS printer is equipped with a 2.3W 455nm blue visible laser. A powder batch of approximately 150 g was used for each build cycle. A CAD file with ten printlet having 5 mm height and 12 mm diameter was loaded onto the Sintratec central software. As mentioned earlier, the print parameters were constant for all print jobs. The layer height, number of perimeters, perimeter offset, hatching offset, hatching spacing was set to 100 μm, 1, 200 μm, 120 μm, and 25 μm, respectively. Furthermore, the processing parameters for the screening conditions and the optimization studies are enlisted in **Table 1** And **Table 2**. For the optimization studies, each manufacturing lot composed of ten printlet, which were tested for their weight, and dimensions using a calibrated weighing balance and a vernier caliper, respectively. Moreover, the tablets from each printed batch were tested for hardness (n=3) (using a TA-XT2 analyzer (Texture Technologies Corp, New York, NY, USA)), disintegration time (n=3), crystallinity, and purity (% degradation). The tablets’ average dimensions were used to calculate the average volume of tablets for each batch using equation 1, where ‘r’ is the radius and ‘h’ is the height of the tablets. The average volume and average weight of each batch were further used to calculate the tablets’ density using equation 2. Density was then used as one of the dependent variables in the DoE for printlet optimization.

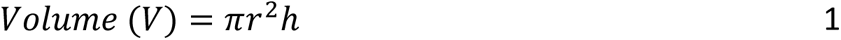

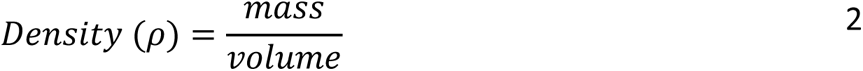

### 2.5 Degradation testing

As a part of the screening studies after sintering the drug-polymer blend, it was imperative to determine the laser’s impact on NFD. For this purpose, high-performance liquid chromatography was used. An analytical technique for the qualitative identification of the degradants was developed for HPLC equipped with a mass spectrophotometer. Moving forward, to quantify the identified degradants, a method for HPLC equipped with a UV-Visible detector was developed.

#### 2.5.1 High-performance liquid chromatography with mass spectroscopy (HPLC-MS)

Samples were analyzed using an Agilent 6530 Q-TOF LC/MS with an Agilent Jet Stream electrospray ionization (ESI) source in positive mode. Chromatographic separations were obtained under gradient conditions using an Agilent Eclipse Plus C18 column (50 x 2.1 mm, 5-micron particle size) with an Agilent Zorbax Eclipse Plus C18 narrow bore guard column (12.5 x 2.1 mm, 5-micron particle size) on an Agilent 1260 Infinity liquid chromatography system. The mobile phase consisted of eluent A (water + 0.1% formic acid) and eluent B (methanol). The gradient was as follows: held at 5% B from 0 to 2 min, 5% B to 20% B from 2 to 5 min, 20% B to 95% B from 5 to 12 min, held at 95% B from 12 to 16 min, 95% B to 5% B from 16 to 16.1 min, and held at 5% B from 16.1 to 20 min. The flow rate was 0.7 mL/min. The sample tray and column compartment were set to 7.5°C and 30°C, respectively. The fragmentor was set to 80 V. Q-TOF data was processed using Agilent MassHunter Qualitative Analysis software.

#### 2.5.2 High-performance liquid chromatography with UV-Visible detector (HPLC-UV/Vis)

The HPLC method from Ma et al. was adapted and modified to better separate the photolytic degradation experienced in the study. Standards were made using methanol while taking precautions to avoid accidental exposure to light. Using a Dionex UltiMate 3000 high-pressure liquid chromatography (HPLC) system (Thermo Scientific, Sunnyvale, CA) equipped with an UltiMate RS Variable Wavelength detector set to 235 nm and Chromeleon 7 software for data acquisition and analysis. During the analysis, the system is held isocratically (70% A: 30% B). The aqueous phase, mobile phase A, consists of HPLC grade water and the organic phase, mobile phase B, consists of acetonitrile. The column separated 10 µL injections with a flow rate of 0.9 mL/min over 30 minutes. A C18, 5 x 20 mm, 5 um columns (Thermo Scientific, Waltham, MA) was used at room temperature to perform the separation.

### 2.6 Printlet Characterization

The crystallinity of the printlet were investigated using X-ray diffraction, and modulated differential scanning (mDSC) analysis, although the mDSC was performed only for the optimized sample. Further, the optimized sample was tested using a pH shift *in vitro* dissolution test to assess the performance of the printlet in comparison to the crystalline drug.

#### 2.6.1 Powder X-ray diffraction studies (PXRD)

A Rigaku MiniFlex 600 (Rigaku, The Woodlands, TX) was utilized to evaluate NFD crystallinity in printed tablets. The instrument is equipped with a Cu-K alpha radiation source. The current is set to 15 mA with a voltage of 40kV. For sample analysis, the printed tablets are crushed into a fine powder, where the powder is evenly spread into an aluminum sample holder and analyzed over a two theta range of 5-40 ° 2θ, a scan speed of 2° per minute, and a step size of 0.02 ° per minute while rotating.

#### 2.6.2 Modulated differential scanning calorimetry (mDSC)

A Q20 DSC unit (TA Instruments, New Castle, DE) conducted modulated differential scanning calorimetry (mDSC) measurements at a heating rate of 3°C/min from 35-200°C. During the experiment, the temperature was modulated by 0.3°C every 50 seconds, with a nitrogen flow of 50 mL/min (Citation of the previous manuscript). For all samples, 8-10 mg was weighed into T-zero pans using a Sartorius 3.6P microbalance (Göttingen, Germany).

#### 2.6.3 Non-sink pH-shift dissolution

A small-volume, non-sink, pH-shift dissolution evaluated the optimized formulation’s solubility enhancement compared to that of the physical mixture. Run 10 floated when placed in the dissolution media and rapidly dissolved by the 10 minute time point. The individual tablet weights for this study were 335.3, 353.5, and 353.6 mg. The weight of the physical mixture used was the same weight of the three tablets, 335.2, 353.7, 353.8 mg. The dissolution media for the study utilized an acidic phase to mimic the stomach, and a neutral phase, to mimic the small intestine. The acidic phase consisted of 0.01 N HCL. The neutral phase consisted of pH 6.8 FaSSIF. A small-volume pH-shift dissolution was performed on an SR8 Plus dissolution tester (Hanson Research Cord., Chatsworth, CA) with 150 mL glass vessels and mini-paddles. A paddle speed of 100 RPM was utilized while the temperature was maintained at 37 °C. The optimized tablets (n=3) and the Physical mixture (n=3) were dropped into 90 mL of 0.01 N HCL. At 30 minutes, 60 mL of FaSSIF (2.24 g/L SIF in 0.1M sodium phosphate buffer) was used for the pH-shift transition to make a total volume of 150 mL. For all sample pulls, 1 mL of the volume was removed and replaced with an equivalent amount of media. Samples were taken at 5, 10, 15, 25, 35, 45, 60, 90, 120, 180 and 240 minutes. All samples were immediately filtered through a 0.22 um PTFE syringe filter and diluted in 1:1 methanol. Caution was taken to avoid light exposure during the dissolution study by covering the apparatus with aluminum foil to avoid accidental light exposure and keeping overhead lights off when not sampling. Sample concentrations were determined by HPLC analysis using the unmodified method previously mentioned by Ma et al.

### 2.7 Dosage form quality assessment (dimensions, microscopy, hardness, and disintegration test)

A VWR® digital caliper (VWR®, PA, U.S.) was used to determine the diameters and thicknesses of the tablets. Images of the printed tablets were taken using Dino-Lite optical microscopy. A texture analyzer (TA-XT2 analyzer, Texture Technologies Corp, New York, USA) along with a one-inch cylinder probe apparatus was used to assess the hardness of the printlet. The test speed was set at 0.3 mm/s and the samples were positioned between the probe across their diameter. The samples’ dimensions were inserted in the software before the test, and the probe stopped at a distance of 3 mm from the starting point of the test, which was deemed sufficient to assess the hardness of the samples. The first point of drop-in force (peak force) was recorded as the hardness of the samples and the test was performed in triplicates. The average hardness of each sample was inserted in the DoE to further assess the impact of the independent parameters on the hardness of the tablets. For the disintegration test, a basket-rack assembly filled with 900mL pH 2 HCl-KCl and maintained at 37±2°C in a 1000 mL vessel was used. Three tablets were placed in the baskets of the oscillating apparatus, operating at a frequency of 29-32 cycles a minute. The timer was started at the beginning of the test and stopped when the tablets were disintegrated completely with no trace of the samples were observed in the basket. The average disintegration time for each run was recorded and reported as a response parameter in the DoE.

## 3 Results

### 3.1 Laser sintering of NFD promotes photodegradation and amorphous conversion

The first hypothesis of this study was that light-sensitive drugs, absorbing visible radiation at any capacity, will interact with the laser during the SLS process. It was further theorized that if the drug interacted with the laser used in the process, it will undergo photo-lytic degradation, and state transformation (melting). A UV-visible absorption analysis was conducted for NFD in both solid and liquid states to demonstrate the drug’s ability to absorb visible radiation. It was observed that NFD, when dissolved in methanol, exhibited considerable absorbance in the visible region (>380 nm). Furthermore, this absorbance in the visible region was also found to be linear, as seen in **Figure 2-B**, i.e., the absorbance increased with an increase in the concentration of NFD in methanol. This phenomenon described by Lambert-Beer’s law has been used in pharmaceutical analysis and for the quantification of drug substances absorbing electromagnetic radiation. Although one limitation of the law is that the linearity fails at higher drug concentration in a solution, where the transmitted radiation is quantified, and absorbed radiation is determined^27^. Hence, for the solid crystalline NFD sample, a reflectance probe was used. The solid samples’ analysis was qualitative to determine the absorbance spectra of solid NFD samples. It was observed that NFD also absorbed radiation at a wavelength corresponding to that of the laser used in the SLS processing i.e., 455 nm as seen in **Figure 2-A**.

**Figure 2.**
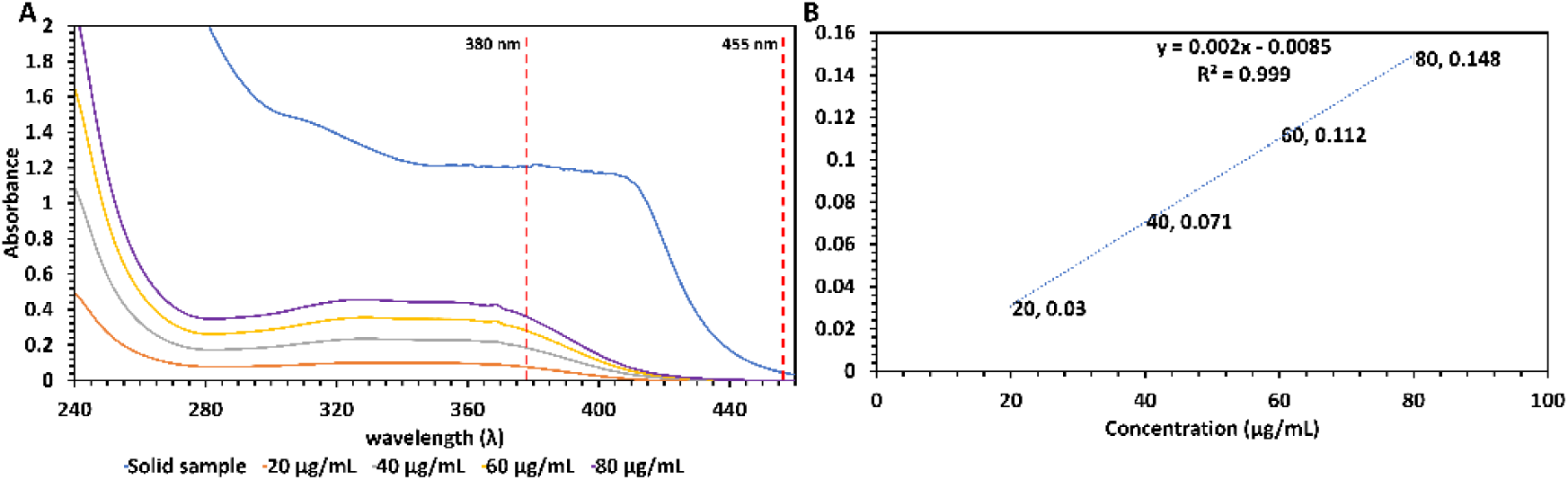
UV-Visible screening studies **A)** UV-Visible spectrum of liquid and solid samples from 460-240 nm wavelength (λ) **B)** increasing absorption with increasing concentration at 400 nm.

From the UV-Visible experiments, it was confirmed that NFD absorbs radiation in the visible spectrum; the next step was to observe the laser’s impact on NFD post SLS processing. For this purpose, the NFD and Kollidon® VA64 physical mixtures (Formulation S1-S3) without a sintering agent were exposed to the SLS process at three different laser speeds (**Table 1**). After the process, the printlets were collected, and their morphology was investigated using microscopy to assess if the parts sintered. The physical evaluation and microscopy indicated that the formulations were sintered in the absence of the sintering agent. This can be attributed to the visible radiation-absorbing ability of NFD. The laser power was sufficient for the drug to absorb radiation and undergo solid-liquid-solid state transformation i.e., melting and solidification, which was confirmed by the amorphous nature of NFD in the printlet post-processing (**Figure 3**). This melting phenomenon can be attributed to the laser absorption because NFD has a T_m_ of 173±2°C, and the surface temperature was maintained at 105°C for these formulations, which is significantly below the melting point of NFD and could not have affected the state of the drug.

**Figure 3.**
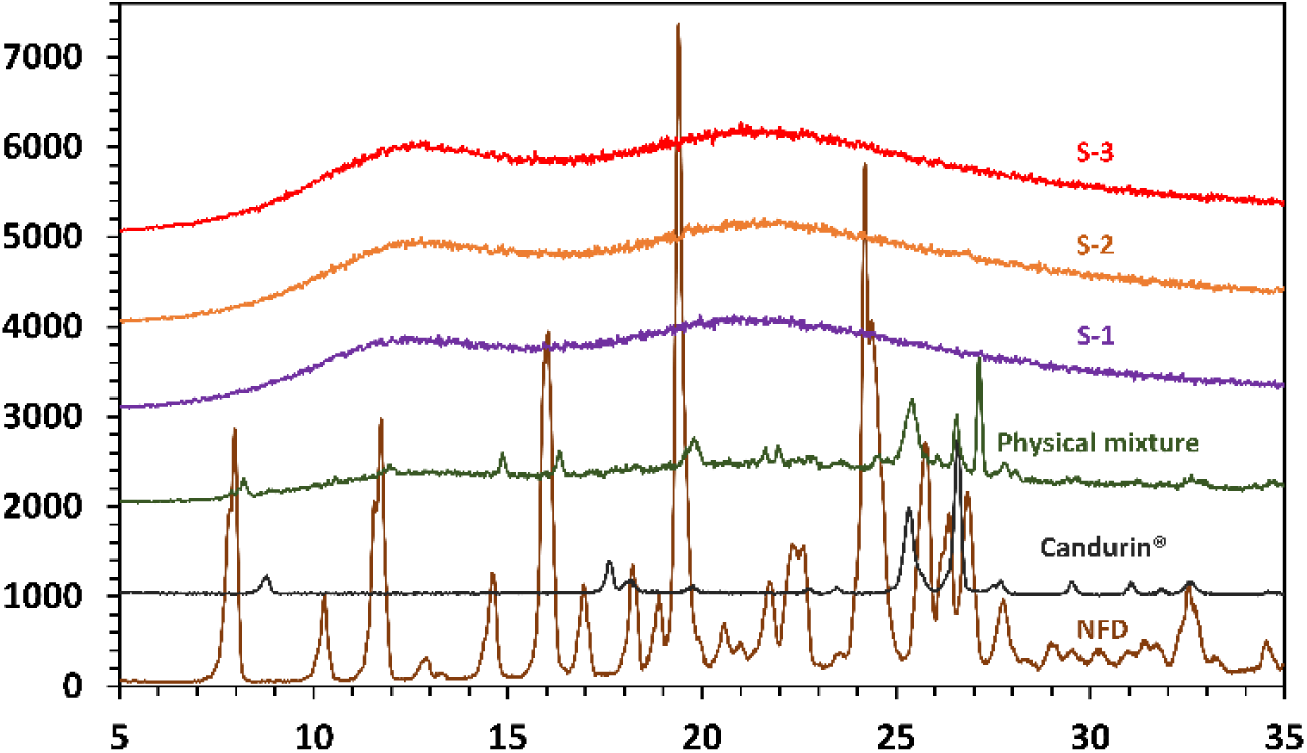
Powder X-ray diffraction spectroscopy of screening samples S1-S3 (10% NFD+90% Kollidon® VA64), Physical mixture for screening samples (NFD+Candurin®+Kollidon® VA64), and pure NFD and Candurin® samples. Kollidon® VA 64 was not included as it is known to be amorphous.

Further, the printlets (Formulation S1-S3) were predominately degraded upon HPLC analysis (e.g., 92.65% nifedipine degradation), **Table 3**. HPLC identified two major degradation products (i.e., Peak 4 and Peak 5) and three minor degradation products (i.e., Peaks 1-3). Therefore, nifedipine’s degradation mechanism was investigated to make the appropriate formulation and process parameter modifications to minimize degradation.

**Table 3.**
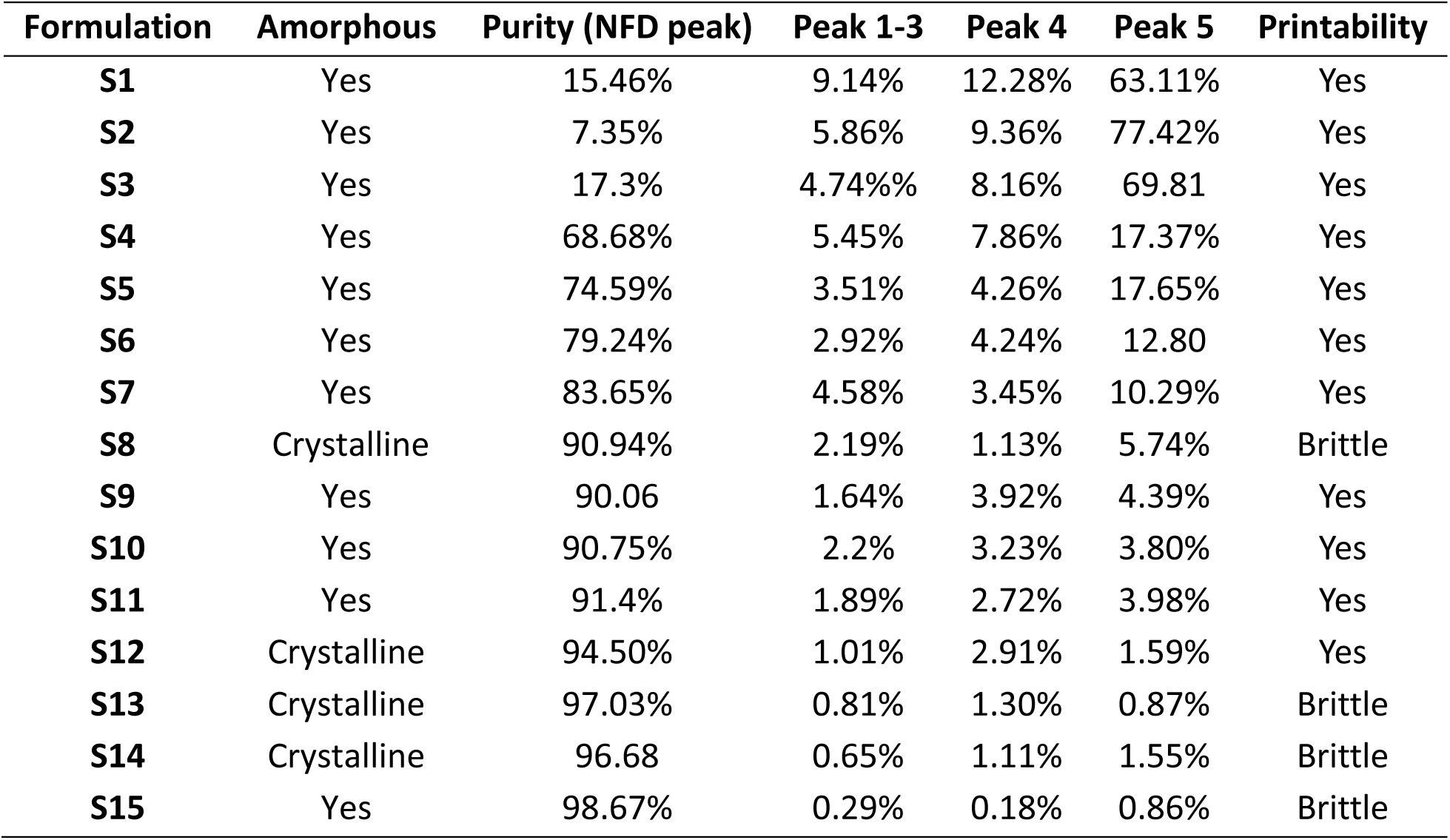
Printability and degradation observations for the screening formulations.

HPLC-MS studies revealed the molecular composition of the two major degradation products (i.e., peak 4 and peak 5). The molecular structure of the degradation products was determined using the molecular composition and the corresponding double-bond equivalents, **Figure 4**. Degradation product 4 results from photolytic degradation caused by visible irradiation of nifedipine; degradation product 5 is from the UV-light mediated oxidation of NFD. For formulations S1-S3, degradation products 4 and 5 contribute to more than 70% of the degradation present. The other minor degradation products (i.e., degradation products 1-3) present during HPLC were not detected by LC-MS as they may be nonionizable species; however, it has been reported that other minor degradation products form during photolytic degradation from inter-molecular interactions amongst nifedipine and the intermediates formed^24^.

**Figure 4.**
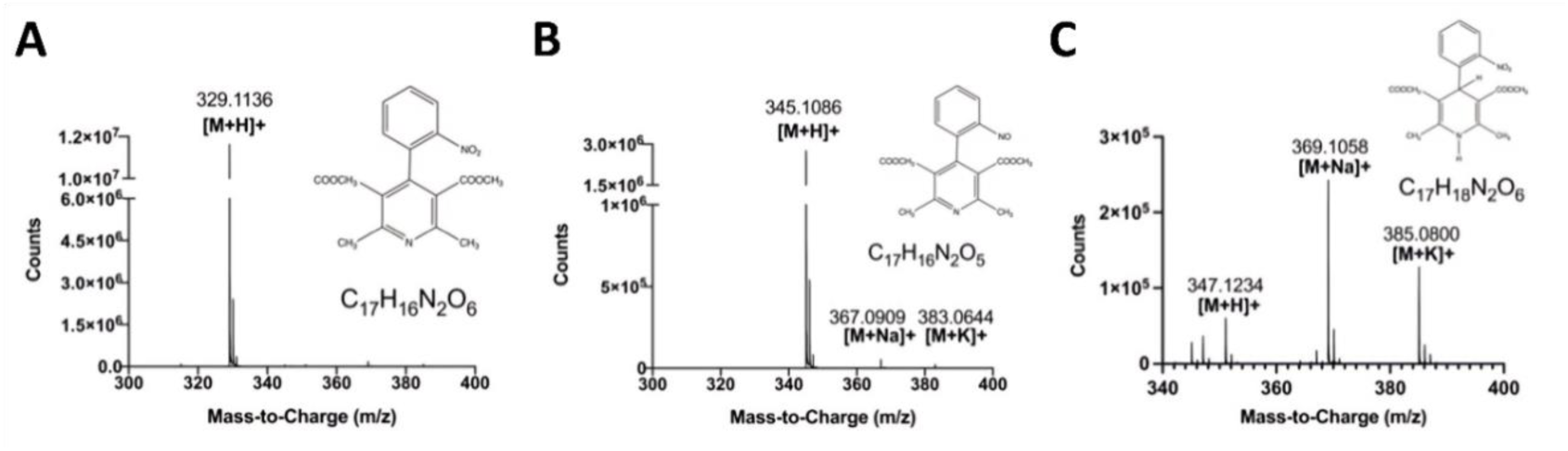
High-performance liquid chromatography-Mass spectroscopy isolated and identified **A)** nitro derivative-oxidative degradation product (UV exposure) **B)** nitroso derivative-photolytic degradation product (visible-light exposure) **(C)** nifedipine.

### 3.2 Screening parameters and range selection

Screening formulations S4-S8 were prepared with a 1:1 ratio (wt%) of NFD and Candurin®. The degradation of NFD in the presence of Candurin® reduced significantly in formulation S4. The difference can be observed in **Table 3**, albeit there still was a considerable amount of degradation (≈32%) present in the printlet at a laser speed of 50 mm/s. Before increasing the amount of Candurin® in the formulation, the laser’s speed was increased to reduce the time NFD was exposed to the laser source. Increasing the laser speed further reduced the degradation from ≈32% to 26%, 21%, 17%, and finally 10% in formulation S5-S8, respectively, where the laser speed was increased from 50 mm/s (Formulation S4) to 250 mm/s (Formulation S8) at a 50 mm/s increment per formulation. The laser speed was not increased any further as the printlets were brittle and exhibited trace crystallinity by PXRD analysis. Formulations S4-S8 provided valuable information about two of the critical parameters in this study i.e., presence of Candurin®, and laser speed, where both impacted NFD’s degradation, and laser speed also influenced the amorphous conversion.

For further parameter screening the drug-to-Candurin® ratio (wt%) was modified, formulations S9-S11 were prepared with a 1:1.5 NFD and Candurin® ratio (wt%). Formulation S9 was processed at a laser speed of 200 mm/s causing 10% degradation, confirming the continued benefit of Candurin® in the formulation. In comparison, formulation S7, at the same laser speed, observed about 17% degradation. Using a 1:1.5 ratio (wt%) of NFD to Candurin®, the surface temperature for formulation S10 and S11 was increased from 105°C to 110°C and the chamber temperature was increased from 80°C to 90°C as under the previous temperature conditions formulation S8 was not printable at 250 mm/s. Although formulations S10 (250 mm/s) and S11 (300 mm/s) were printable on increasing the surface temperature, the change in degradation with increasing laser speed was not significant as seen in **Table 3**. These results point out the impact of surface temperature on the degradation and amorphous conversion of NFD, which was previously not predicted. Hence surface temperature was also deemed as a critical parameter for this study.

Moving forward, the ratio (wt%) of NFD to Candurin® was increased to 1:3 for formulations S12-S15. The degradation observed for formulation S12 was about 5%, which was significantly less compared to S11, which was about 9%. Even though formulation S12 was printable, it was found to have trace crystallinity, and on further increasing the laser speed to 400 mm/s, it was brittle and had about 3% degradation. On increasing the surface temperature to 115°C, 4% degradation was observed with a brittle printlet and trace crystallinity. On further increasing the surface temperature to 120°C and the laser speed to 450 mm/s, <2% of degradation was observed along with complete amorphous conversion, however, the printlet was found to be brittle.

From the results of these screening studies, it was evident that the level of Candurin®, laser speed, and surface temperature play a critical role in the degradation of NFD. Moreover, laser speed and surface temperature also play a role in the amorphous conversion and printability of the printlet. Hence these three parameters were considered as independent variables for the DoE. Moreover, from the screening studies, the printable range for each of the parameters were selected where Candurin® was used at 1:1, 1:1.5, and 1:3 ratios (wt%) with the drug, the laser speed was set with a minimum of 200 mm/s and a maximum of 300 mm/s, and the surface temperature was set at a minimum of 100°C and maximum of 120°C.

### 3.3 Optimization studies

After manufacturing all the formulation compositions using different processing conditions, the manufactured printlets were subjected to various characterization techniques. The data collected from the experiments was introduced as responses to the DoE. **Table 4** is a collection of numeric values inserted into the DoE to understand the relationship between each independent variable (Candurin®, laser speed, and surface temperature) on the response variables (crystallinity, purity, hardness, weight, density, disintegration time), which is discussed in-depth in the following sub-sections.

**Table 4.**
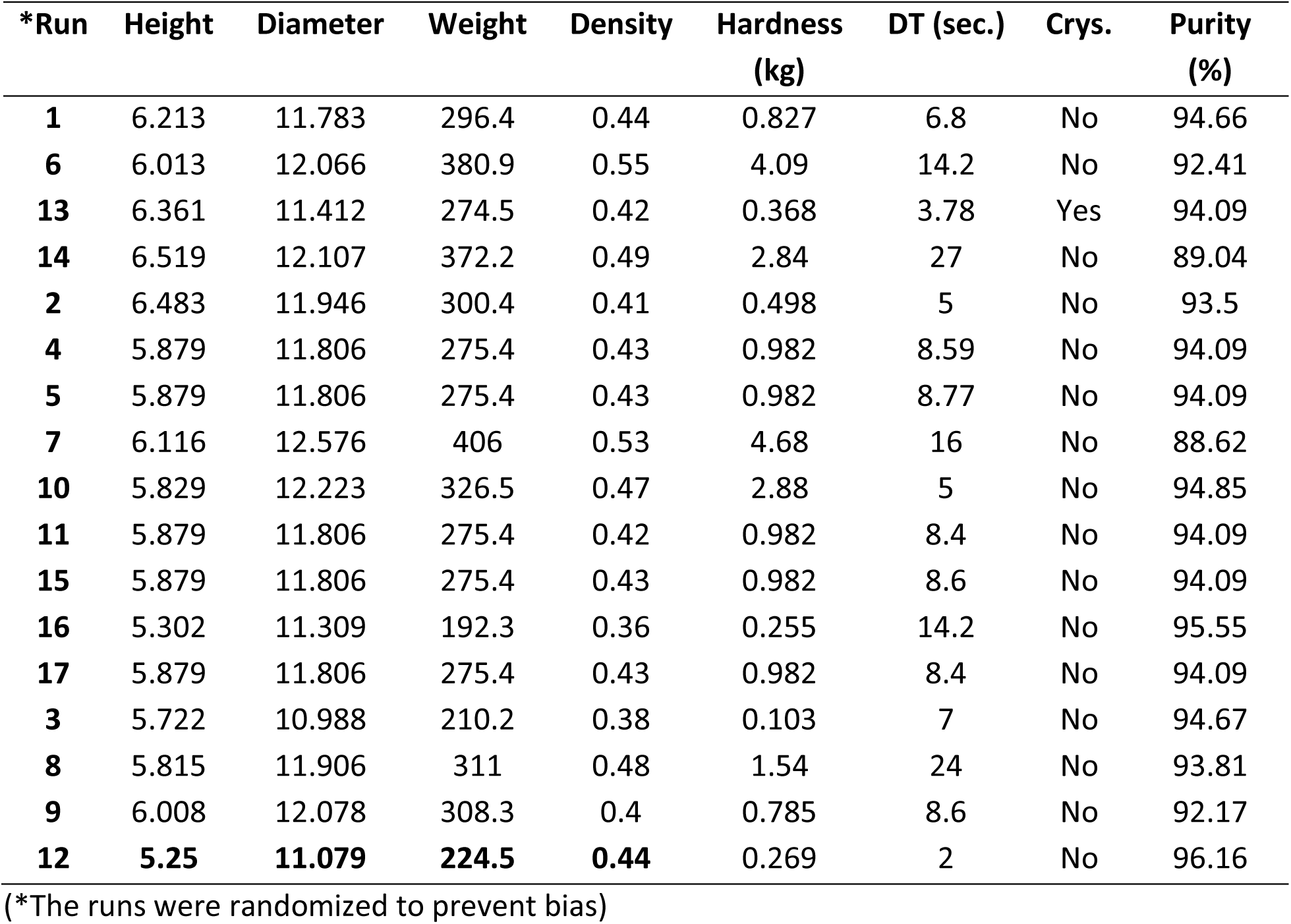
Compilation of experimental responses for different combinations of independent variables (Runs 1-17).

#### 3.3.1 Crystallinity

PXRD was used to determine the crystallinity of NFD in the DoE formulation. From the screening studies, increasing the laser speed led to crystallinity or partial amorphous conversion in the formulation. For the DoE samples, the laser speed was maintained at or below 300 mm/s; thereby, it was expected that all the formulations will undergo amorphous conversion and subsequent formation of an amorphous solid dispersion. From the XRD results depicted in **Figure 5**, all samples, except for Run 13, demonstrated the absence of crystalline peaks. The two-theta (2θ) values for these experiments were set from 20-30 degrees as the physical mixture demonstrated strong NFD crystalline peaks in this region.

**Figure 5.**
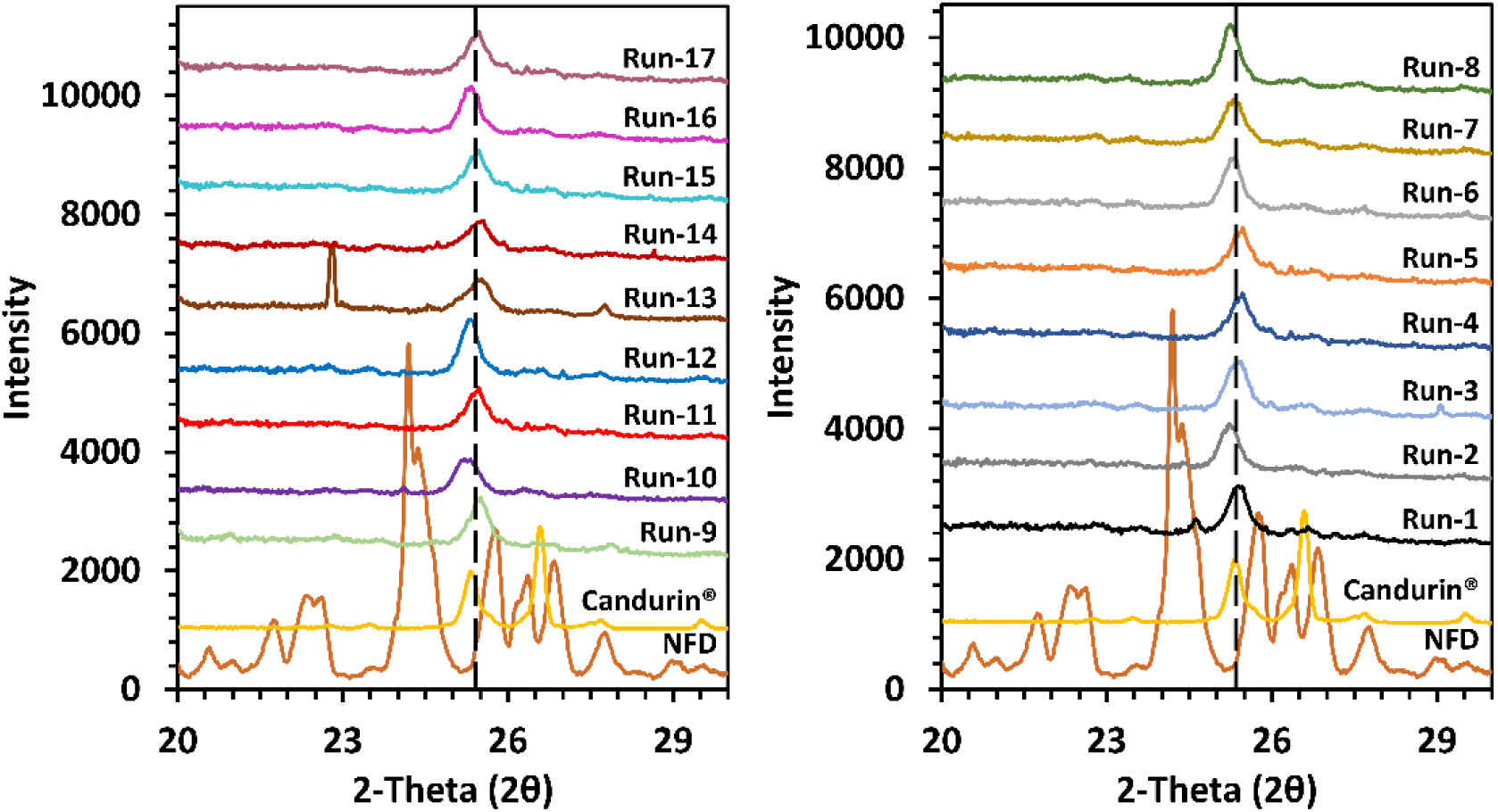
Powder X-ray diffraction spectroscopy of DoE samples (Run 1-17), The two-theta (2θ) values from 20-30 were selected based on the crystalline peaks observed in the physical mixture in Figure 3. The broken lines represent Candurin® peak at a 2θ value of 25 degrees.

Moreover, due to the presence of Candurin®, which demonstrated 2θ values at 8.9, 17.6, 18.4, 25.3, and 26.5 degrees, it was also included in the overlay created for the analysis. Characteristic NFD peaks can be seen in **Figure 5** at 2θ values of 22.4, 24.1, 25.7, 26.6 degrees. The NFD peaks are absent in all the DoE samples except for Run 13, which consisted of 5% Candurin® and was manufactured at a laser speed of 250 mm/s with a surface temperature of 100°C. This may be attributed to the low surface temperature maintained for manufacturing the printlet. In the screening experiments, we observed a relationship between surface temperature and amorphous conversion, where an increase in surface temperature facilitated amorphous conversion as a function of higher energy input. Surface temperature’s impact on amorphous conversion was confirmed by observing Run 1 and Run 14, which have similar compositions as Run 13 but were manufactured at a higher surface temperature (110°C) and Run 1 was processed at a faster laser speed (300 mm/s) than Run 13. Run 3, Run 16 and Run 2 were also manufactured at a surface temperature of 100°C, although they observed complete amorphous conversion. Amongst these runs, Run 3 was processed at the same manufacturing conditions as Run 13 but contained 15% w/w Candurin®. This comparison is interesting as it suggests that Candurin® also plays a role in amorphous conversion and increasing the amount of Candurin® in the formulations facilitates the amorphous conversion of crystalline NFD. Candurin® facilitating amorphous conversion is also seen in Run 16 and Run 2, which have higher amounts (10% w/w) of Candurin® as compared to Run 13 (5% w/w). The peaks which are consistent in all formulations at a 2θ value of 25.3 degrees correspond to the Candurin® peaks and should not be mistaken as the presence of crystallinity in the runs.

#### 3.3.2 Degradation

Laser-induced degradation was the key aspect and parameter for this study. From the screening experiments, the SLS process led to extensive degradation of NFD when no photo-absorbing species, such as Candurin®, were used. It was observed that increasing the ratio (wt%) of Candurin® to NFD reduced the degradation observed in the printlet. Moreover, the screening studies observed the influence of laser speed and surface temperature on NFD degradation, which required further assessment.

Box-Behnken is a frequently used Response Surface Methodology based second-order design alongside 3^k^ factorial and central composite designs^28–30^. Box-Behnken has the advantage of not including all the combinations in which all variables are on the highest or the lowest levels^31–33^. This reduces the number of runs while maintaining the integrity of the design. Moreover, for such optimization studies, preliminary screening experiments to narrow down the minimum and maximum values of the variables is imperative, which was conducted in this study. The use of the Box–Behnken design is popular in industrial research because it is an economical design and requires only three levels for each factor where the settings are −1, 0, 1 (see **Figure 1**)^30^.

It was observed that multiple interactions occurred between the response and the independent variables after adding responses to the design points; hence the design was fit into a quadratic model. The model was observed to have an F-value of 34.62, which implies the model is significant, and there is only a 0.01% chance that an F-value this large is due to noise. It was also observed that the individual variables, i.e., Candurin® (F=47.43, p=0.0002), surface temperature (F=39.95, p=0.004), and laser speed (F=182.82, p=<0.0001) demonstrated a significant impact on the degradation of NFD. The impact of these variables was not only significant, but they also demonstrated a correlation with the degradation, which can be seen in **Figure 6 (A, B, C)**. The trend that was observed indicates that an increase in the ratio (wt%) of Candurin® to NFD, and an increased laser speed reduce the degradation caused by the process (increase the purity), whereas an increase in surface temperature reduces the purity and increases the degradation observed. This confirms the assumptions made for the laser speed and surface temperature while analyzing the screening formulations.

**Figure 6.**
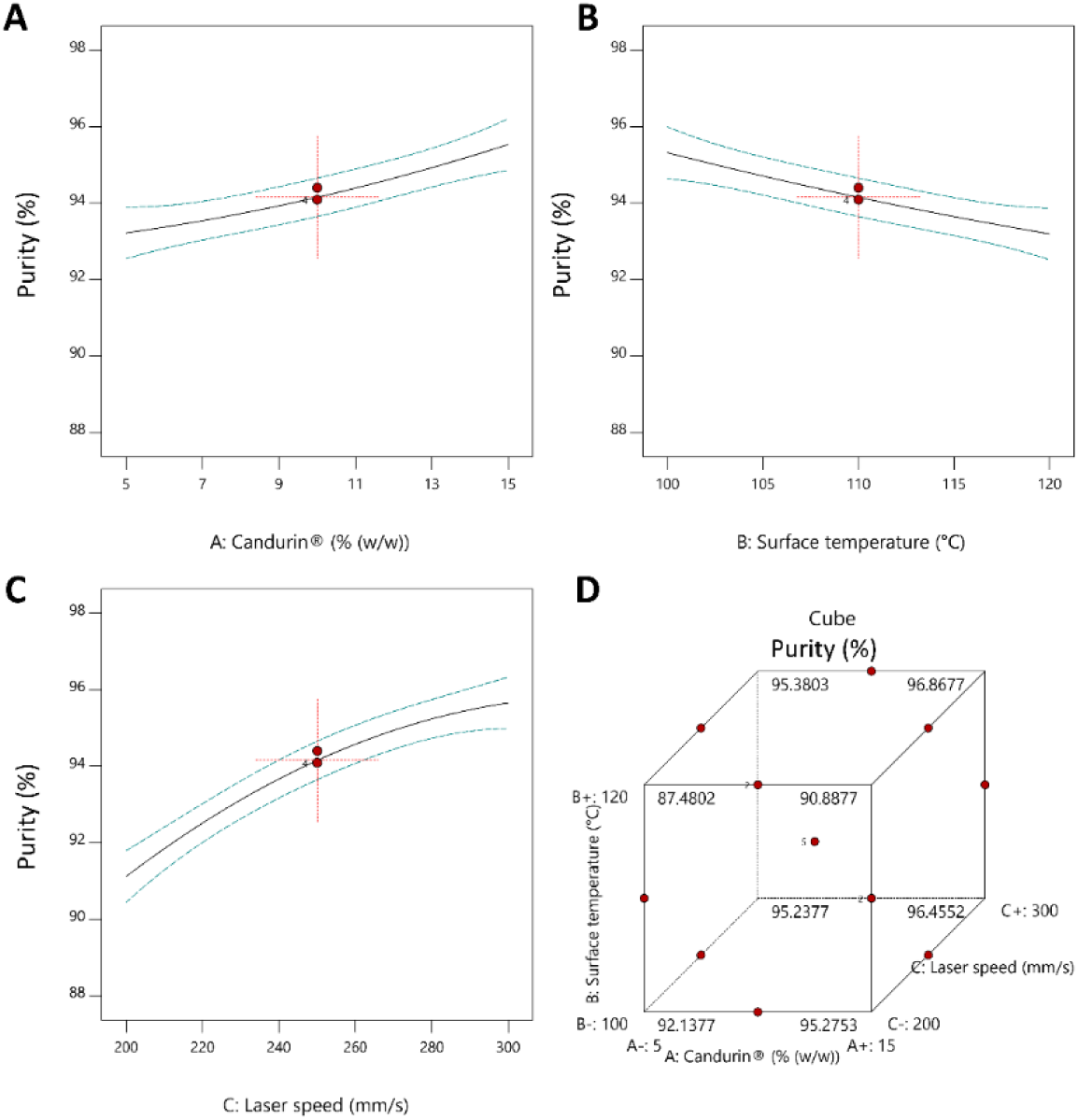
Variable-response relationship trends between % Purity and **A)** Candurin® (wt%), **B)** Surface temperature (°C), **C)** Laser speed (mm/s), **D)** All three independent variables.

Furthermore, it was also determined that a combination and interplay between the two processing variables, i.e., surface temperature and laser speed, had a significant impact (F=25.54, p=0.0015) on the purity of the samples. The model suggested that laser speed observed the most significant impact on the degradation of NFD during processing amongst all the independent variables. Laser speed’s impact can be observed in **Figure 6 (D),** where the highest purity values correspond to the axes with the highest laser speed i.e., 300 mm/s.

For formulation and process optimization, one key parameter is the design’s ability to accurately predict change in response to changing a studied variable. This ability can be determined by the ‘Adeq Precision’ of the model, which measures the signal-to-noise ratio^34, 35^. For this model, a ratio greater than 4 is desirable, and for this design, it was found to be 21.069, which indicates an adequate signal and that this model can be used to navigate the design space. Coefficient estimates or contour lines (**Figure 7**) can be used to navigate within the design space. The coefficient estimate represents the expected change in response per unit change in factor value when all remaining factors are held constant. For the tested variables i.e., Candurin®, surface temperature, and Laser speed, the coefficient estimates were found to be 1.16, −1.06, and 2.27 units, respectively. The negative coefficient represents the inverse correlation between surface temperature and purity i.e., purity reduces on increasing surface temperature.

**Figure 7.**
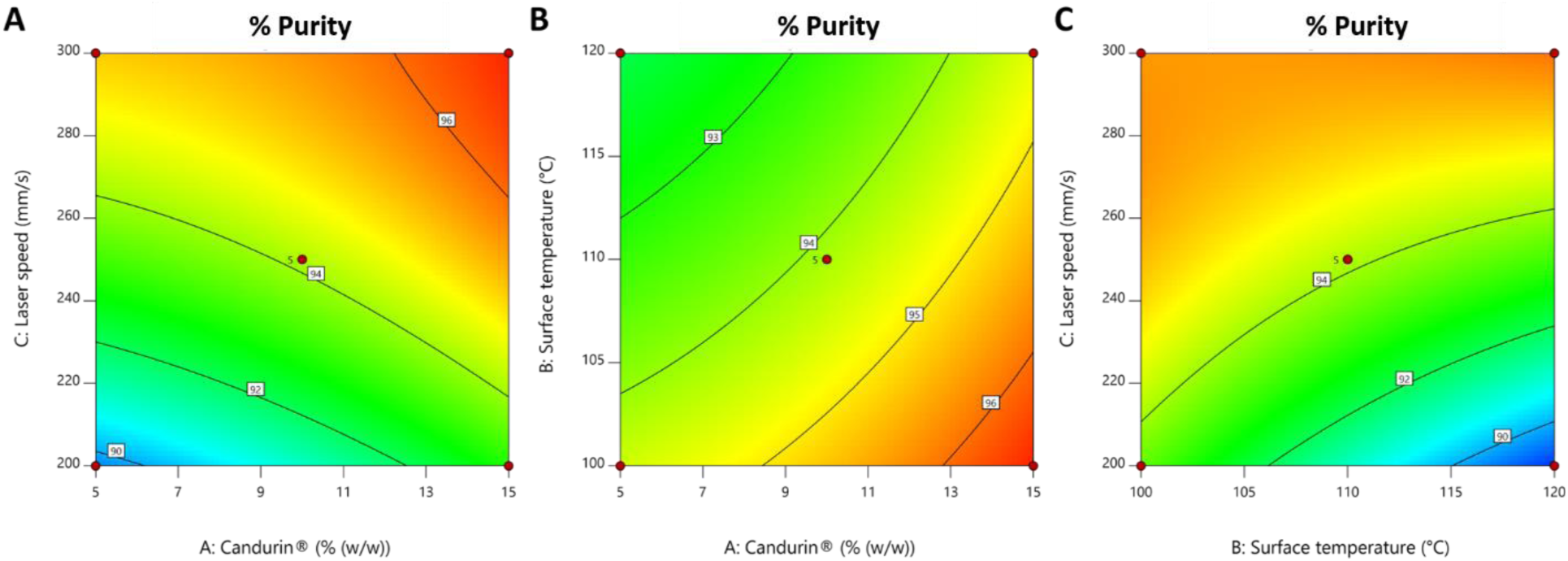
Contour lines representing constant values of % Purity over variable values of **A)** Laser speed and Candurin® **B)** Surface temperature and Candurin® **C)** Laser speed and Surface temperature.

#### 3.3.3 Quality attributes (Hardness, density, weight variation, and disintegration time)

In our previous study, we observed that different processing parameters demonstrated variability in weight, dimensions, and tensile strength of the printlet. In the previous study, assessing the correlation between the processing parameters and these quality attributes was beyond that study’s scope^14^. The current 17-Run study provided an opportunity to investigate the impact of print speed and surface temperature, along with the formulation composition on these critical quality attributes.

##### 3.3.3.1 Hardness

The response values for hardness ranged from 0.013 to 4.68 kg/mm^2^ leading to a maximum to minimum response ratio of 45.44. A ratio of more than 10 indicates that a transformation is required; therefore, a square root transformation was performed. The same quadratic model was used because of interactions between independent variables and their impact on the response, as explained in the previous section. The overall model was found to be significant (F=81.95, p=<0.00001). In this case Candurin® (F=104.76, p=<0.0001), surface temperature (F=511.09, p=<0.0001), laser speed (F=67.38, p=<0.0001), Candurin®-Surface temperature (F=10.11, p=0.015) and, Candurin®-laser speed (F=6.86, p=0.03), were found to be significant. The signal-to-noise ratio (32.062) indicated that this model can be used to navigate the design space. The coefficient estimates for all the significant terms, i.e., Candurin®, surface temperature, laser speed, Candurin®-Surface temperature and, Candurin®-laser speed, were −0.2121, 0.6232, −0.2263, −0.1239, and 0.1021 units, respectively. These coefficients indicate that Candurin® and speed have a negative correlation to the hardness of the printlet. This correlation can be seen in **Figure 8 (A, B, and C),** where an increase in the amount of Candurin® reduces the hardness, and laser speed reduces the hardness of the printlet. In contrast, an increase in the surface temperature increases the hardness of the printlet, which is seen along the axes of the highest value of surface temperature (120°C) in **Figure 8 (D and E)**.

**Figure 8.**
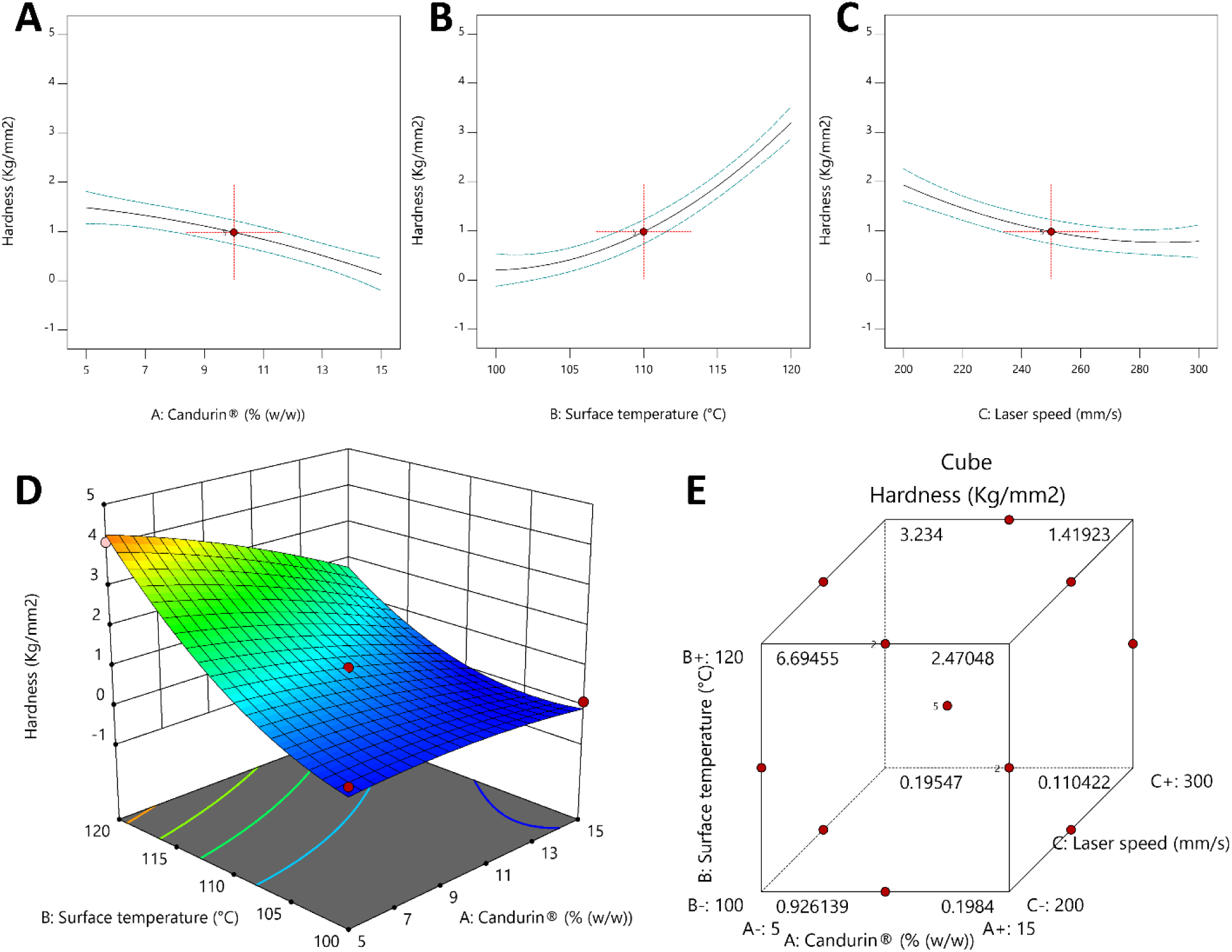
Variable-response relationship trends between hardness and **A)** Candurin® **B)** Surface temperature **C)** Laser speed **D)** 3D surface plot for all three variables **E)** Variable-response cube for all three variables.

Moreover, the complex interactions between different independent variables on hardness can be observed in the 3D surface plot in **Figure 8**. This observed relationship can be explained by the change in the formulation composition from the increase in Candurin® i.e., the amount of Kollidon® VA64 reduces. Candurin® is merely a sintering agent, the sintering occurs due to the thermoplastic nature of Kollidon® VA64 as it absorbs the heat conducted by the sintering agent, undergoes thermal transition, and solidifies, resulting in the sintering of nearby particles together. This data demonstrates that an increase in Candurin® (reduction in Kollidon® VA64) reduces the process’s sintering efficacy and leads to brittle structures with low tensile strengths. To see this practically, a direct comparison can be made between Run 6 and Run 8, which are processed at the same conditions (surface temperature: 120°C, laser speed: 250 mm/s), but the former has 5% Candurin® (90% Kollidon® VA64) and latter has 15% Candurin® (80% Kollidon® VA64). Run 6 was found to have a hardness of 4.09 kg/mm^2^, whereas Run 8 had a hardness of 1.54 kg/mm^2^. Further, Run 6 (120°C) can also be compared to Run 13 (100°C) to show the impact of surface temperature with other variables constant on the hardness, where Run 13 observed a hardness of 0.368 kg/mm^2^. Additionally, Run 1 (300 mm/s) and Run 14 (200 mm/s) can be used to demonstrate the impact of laser speed when both formulations were processed at 110°C surface temperature with 5% Candurin® in their formulation and demonstrated a hardness of 0.83 kg/mm^2^ and 2.84 kg/mm^2,^ respectively.

##### 3.3.3.2 Density and weight variation

Weight variability resonates closely to drug content uniformity and dose of the printlets, whereas density relates the dimensions of the printlet to the weight^36^; hence these two response variables were considered for the evaluation of quality attributes. For both weight and density, the maximum to minimum response ratio was below 10, and hence no transformations were conducted. The data was fit into a quadratic model similar to the above sections. Both weight (F=174.50, p=<0.0001) and density (F=33.80, p=<0.0001) models were found to be significant. All independent variables (Candurin®, surface temperature, laser speed) were found to have a significant impact on the weight (F=275.94, p=<0.0001; F=756.31m p=<0.0001; F=456.29, p=<0.0001) and density (F=25.30, p=0.0015; F=215.83, p=<0.0001; F=29.86, p=0.0009) of the printlet. The surface temperature-laser speed significantly impacted the weight of the printlet (F=6.19, p=0.047), whereas Candurin®-laser speed (F=6.57, p=0.0374) had a significant impact on the density of the printlet. A signal-to-noise ratio of 45.049 and 20.805 was detected for the weight and density responses, suggesting that this model can be used to navigate the design space. Candurin® and laser speed were found to negatively correlate with the weight and the density of the printlet; their coefficients were found to be −33.75, and −43.40 units for weight and -0.02 units for both the variables for density of the printlet. The significance of the coefficients has been explained in the previous sections. The coefficients for surface temperature were positive for both weight (55.88 units) and density (0.058 units), which means an increase in surface temperature increases the tablets’ weight and density. These relationships can be observed from the cube and 3D surface plots for weight and density in **Figure 9**. The reason behind this trend can be explained by the sintering phenomenon, where a slower laser speed at a higher surface temperature dissipates more energy on the surface as compared to a higher laser speed at a lower surface temperature. This energy causes the thermal conversion of the polymer, leading to an increase in the density of the layer and a reduction in the porosity, which forms a cavity on the print surface during the printing process. The higher the energy dissipation, the steeper the cavity. When the next layer of powder is spread onto this surface, more powder gets filled in the steeper cavity, which gets sintered by the laser, this is responsible for a larger weight of the tablets even though the print dimensions are the same in both these cases.

**Figure 9.**
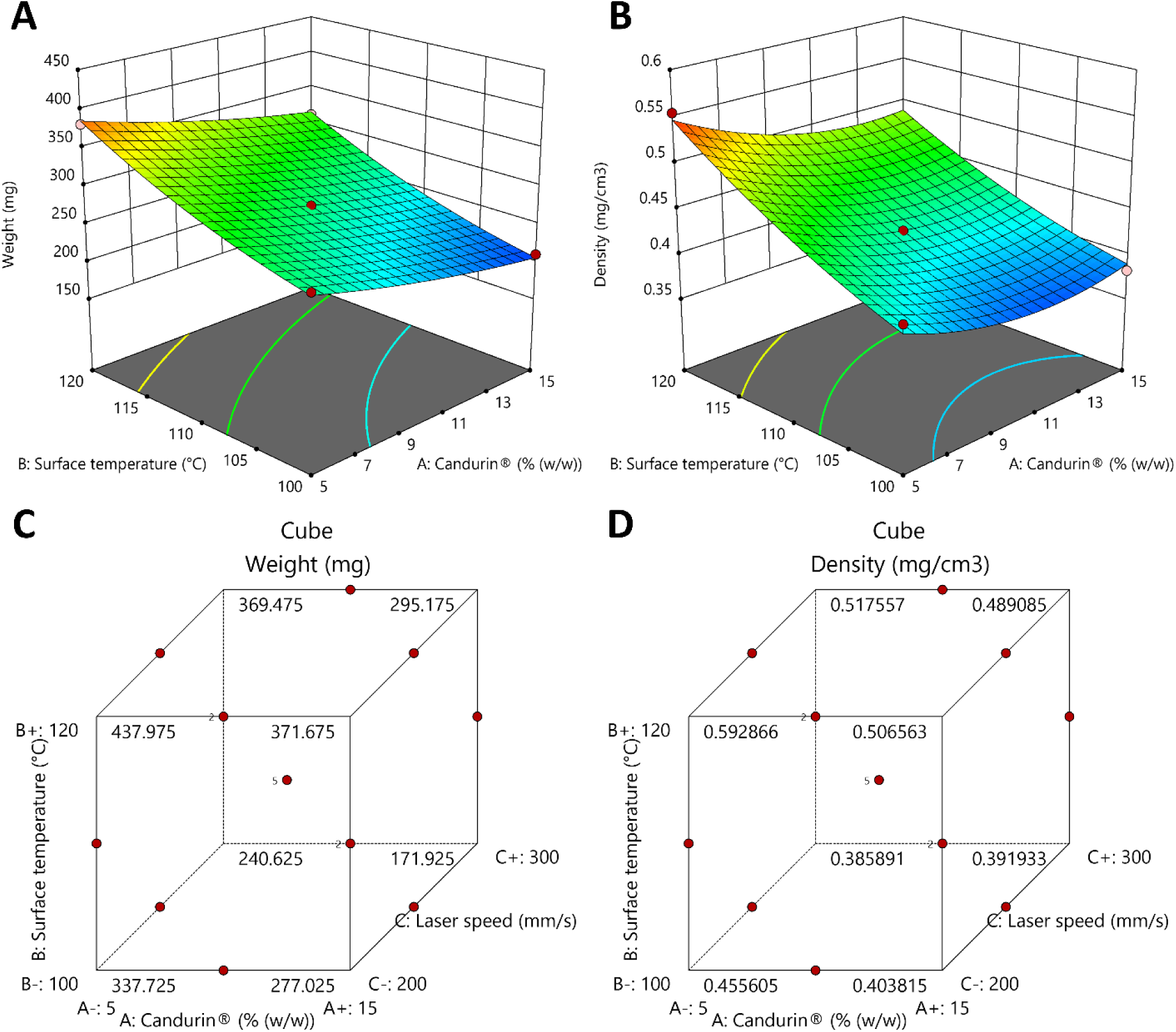
3D response surface plot for **A)** Printlet weight against all three variables **B)** Printlet density against all three variables. Variable-response cube for **C)** Printlet weight against all three variables **D)** Printlet density against all three variables.

**Figure 10.**
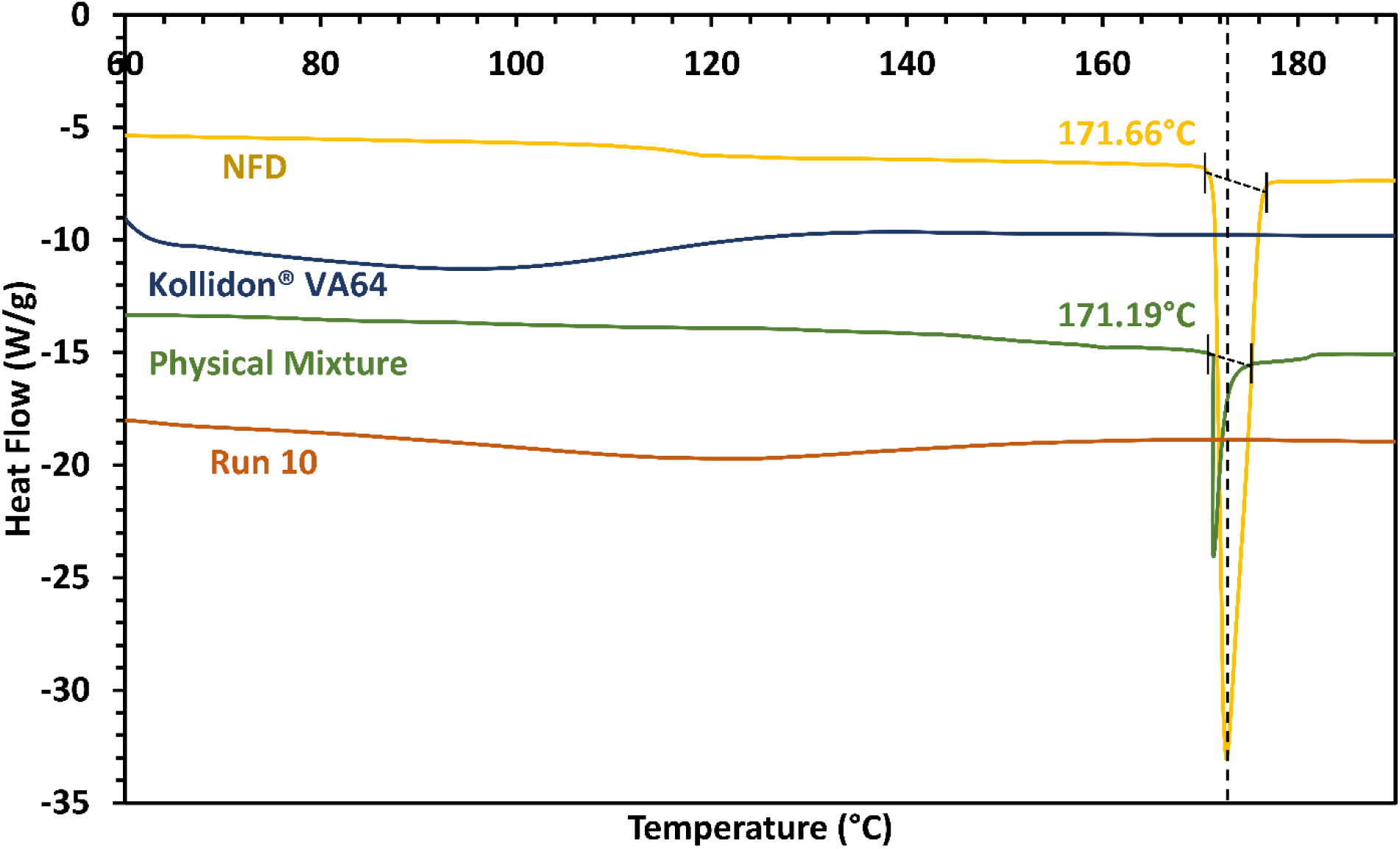
Differential scanning calorimetry to confirm amorphous conversion in the optimized formulation.

This can be practically seen in Run 7 where the surface temperature is the maximum (120°C) and the laser speed set to a minimum (200 mm/s), resulting in a total weight of ≈406 mg, which is the maximum observed weight in this design. Run 7 can be compared with Run 16, which observes a weight of ≈192 mg where the surface temperature is maintained at the minimum value and the laser speed at the maximum value for this design i.e., 100°C and 300 mm/s. In both these cases, the amount of Candurin® was constant, i.e., 10%. The impact of Candurin® on the weight of the tablets can be assessed by observing Run 6 (5%) and Run 8 (15%) where all the other variables are kept constant (120°C, and 250 mm/s). Run 6 was found to weight ≈380 mg, whereas Run 8 weighted ≈311 mg. This relates to the previously discussed impact of Kollidon® VA64 on the formulation, where the thermal transition of the polymer can increase the hardness of the tablets, which can, in turn, be related to the density of the tablets. All these findings can be used to determine the processing condition and set dimensions in the CAD model for manufacturing dosage forms with a target weight. These trends also help understand the interplay between the processing parameters and the formulation parameters in an SLS 3D printing process.

### 3.4 Characterization of the optimized formulation

Though degradation is the key aspect and parameter of this study, the formulations with the lowest degradation levels (i.e., Run 12 and Run 16) were not selected for characterization, as they did not have the best overall printlet characteristics (e.g., hardness). Therefore run 10 was chosen as the optimized formulation for characterization, as these printlets achieved marginally higher degradation (i.e., ∼1%) while having increased printlet hardness. In addition to the characterization reported in **Table 4**, Run 10 was subject to additional characterization to evaluate the amorphous nature of the printlet further; specifically, if the formulation is miscible and provides solubility enhancement through forming an amorphous solid dispersion.

Before evaluating the printlet’s solubility enhancement, the miscibility and amorphous nature of the printlet were further investigated. Upon mDSC analysis, the printlet exhibited a single T_g_ onset at 89 °C; the presence of a single T_g_ suggests a miscible formulation with increased stability. The mDSC data also confirmed the prior PXRD characterization, in that, the formulation did not exhibit any melting endotherms, suggesting the absence of crystallinity. The solubility enhancement of the optimized formulation (i.e., Run 10) was evaluated using a non-sink pH-shift small volume dissolution study. The optimized formulation achieved a quicker and greater extent of NFD release in the acidic phase, achieving a 21-fold and a 3.4-fold increase in solubility compared to the crystalline NFD and physical mixture before the pH transition, respectively (**Figure 11**). Upon pH transition, in the optimized formulation, NFD maintained supersaturation for the study’s duration, achieving a 6.7-fold increase and a 1.8-fold increase in solubility compared to the crystalline NFD and physical mixture at the duration of the study.

**Figure 11.**
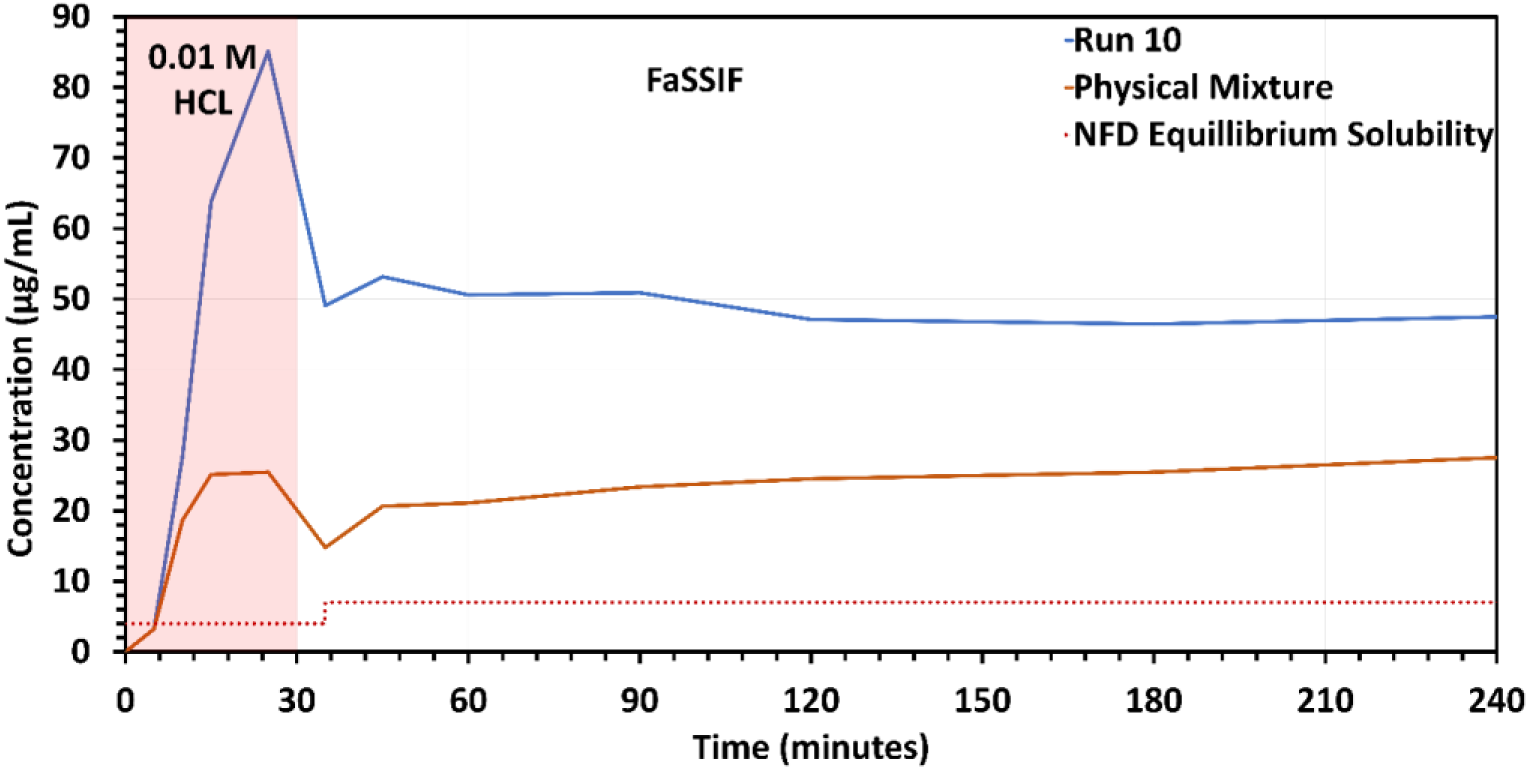
pH shift *in vitro* dissolution testing for Run 10, physical mixture, and crystalline NFD. The change in drug concentration at the 35-minute time point is attributed to the dilution of the dissolution medium from 90 mL to 150 mL.

## 4 Discussion

This study demonstrates the utility of a simple pre-formulation UV-Visible absorption experiment to predict a drug’s ability to act as an electromagnetic energy-absorbing species during SLS. It was also shown that this laser absorbing activity may lead to electromagnetic radiation-mediated degradation and solid-state transformation of the drug. Although the drug degraded under the influence of the process, it still sintered the drug-Polymer physical mixture in the absence of Candurin®. Thereby it can be confirmed that if the drug is stable under the influence of the laser, it can aid the sintering process and reduce the amount or eliminate the need for excipients such as Candurin® in the formulation. In a contrary case where the drug undergoes photolytic degradation, photo absorbing species such as Candurin® that has been used as opacifying agents in the pharmaceutical and cosmeceutical industry can be used to prevent laser mediated degradation. This was demonstrated in the current study for nifedipine, which possesses both π-bonds and non-bonding orbitals (lone pairs in ‘N’ and ‘O’) hence is extremely sensitive to ultraviolet radiation and visible light up to 450 nm. Previous studies have observed that nifedipine gives nitrosophenylpyridine homolog on exposure to daylight, and nitro-phenylpyridine homolog on UV irradiation. This vulnerability towards electromagnetic radiation made NFD an excellent model drug for this study^25^.

In an attempt to overcome the degradation that arises when printing NFD:VA64 powder blends, an understanding of the degradation pathway was required to make educated modifications to the process parameters and formulation composition. Therefore, LC-MS was used to determine the molecular formula of the two degradant products identified during the HPLC analysis of the NFD:VA64 printed tablet. NFD undergoes both photolytic degradation and photo-oxidation^24, 37, 38^. The molecular formula of the degradants detected by LC-MS i.e., C_17_H_16_N_2_O_5_ (2,6-dimethyl-4-(2-nitrosophenyl)-3,5-pyridinedicarboxylic acid dimethyl ester) also known as NTP and C_17_H_16_N_2_O_6_ (2,6-dimethyl-4-(2-nitrophenyl)-3,5-pyridinecarboxylic acid dimethyl ester) also known as oxidized nifedipine, align with previously reported electromagnetic light-mediated degradation products^38, 39^. Nifedipine on irradiation mainly converts to NTP which is a stable paramagnetic species reported by Damian and colleagues (2006)^38^. Furthermore, electron paramagnetic spectroscopy (EPR) revealed that an increase in the irradiation time also increased the intensity of the EPR signal, hence the degradation and radical formation were irradiation time dependent^38^. This helps understand the impact of laser speed on the degradation of NFD. The DoE observed a significant impact of laser speed on the degradation where a higher laser speed (lower exposure time) led to a reduced degradation. Moreover, the kinetics of photo-degradation and photo-oxidation determined by Majeed and colleagues (1987) demonstrated the impact of a variable light source and temperature, where different light sources depicted the different extent of degradation with the highest degradation at 380 nm^39^. These findings help understand the impact of the surface temperature on the degradation of NFD as the DoE observed a significant impact of the surface temperature on the extent of degradation of NFD. An increase in temperature led to reduced purity of NFD in the printlet which can be attributed to the lamp placed over the print surface and used as a heat source for SLS printing^40^. The quantum yield for photodegradation is about 0.5; statistically which means that of every two photons absorbed, one causes decomposition of a nifedipine molecule which led to the almost complete degradation of NFD in formulations without Candurin®, whereas on adding Candurin® it coated the NFD crystals and competed with NFD to absorb the electromagnetic energy^38^. With all other printing parameters unchanged, incorporating Candurin® limited the amount of energy NFD absorbed, and the degradation of NFD was decreased. Moreover, the amount of Candurin® had a significant impact on the purity of NFD in the printlet as shown by the DoE. These findings suggest that SLS processing has limited use for processing light-sensitive drugs at this point, as a combination of high laser speed and low surface temperature along with additional formulation considerations, such as the addition of photo-absorbing, opacifying agents is required.

For this study, the transformation of the drug to its amorphous form was important as NFD is a class II drug as per the biopharmaceutical classification system (BCS) and exhibits dissolution limited absorption, and bioavailability^41, 42^. Such molecules can be formulated as supersaturating drug delivery systems such as amorphous solid dispersions for an increase in solubility and dissolution rate^43^. The optimized formulation in this study was found to have a 21-fold increase in solubility as compared to the crystalline NFD before the pH transition and a 6.7-fold increase in solubility after the pH shift. The solubilized drug remains stable at both pH conditions, this trend agrees with previously conducted studies by Theil and colleagues (2016), and Ma and colleagues (2019) that demonstrate solubility enhancement and stability of NFD in Kollidon® VA at the drug load used in the current study. These findings along with the XRD and DSC observations conclude the formation of an ASD post-SLS processing.

Apart from the purity, crystallinity, and performance of the printlets, other critical quality attributes such as printlet dimensions, tablet weight variation, hardness, and density were also assessed as a part of this study. It was found that the processing and formulation parameters have a critical and significant influence on these parameters, where an increase in the laser speed, amount of Candurin®, and decrease in surface temperature led to a reduced hardness and average weight of the tablet. Fina and colleagues (2018) observed a similar trend between the laser speed and printlet weight, and hardness where they attributed this to higher energy input from the laser leading to more number necks forming in each layer at lower laser speeds and reduced empty spaces providing more room for powder particles to be sintered thereby creating a heavier printlet^18^. However, this relationship was based on observations and only accounted for the impact of the laser which is partially true and can be explained from equation 3:

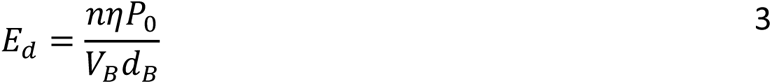

Where ‘*E_d_*’ is the laser power density, ‘n’ are the number of beam passes, ‘*η’* is the absorptivity of the material ‘*P_0_*’ (W) is the beam power, ‘*V_B_*’ (mm/s) is scanning speed and ‘*d_B_*’ (mm) is the beam spot diameter^44^. The equation suggests that the laser power density is inversely proportional to the laser scanning speed and agrees with the observations made by Fina and colleagues (2018). However, the equation does not account for the contribution of surface temperature set on the total energy the surface is exposed to. As per our observations in an SLS process, the heat source exposes the surface of the powder bed to a baseline thermal energy which depends on the set surface temperature, hence the total energy the surface is exposed to is also attributed to the baseline energy from the heat source not just the energy induced by the laser. This was observed in the DoE where an increase in surface temperature led to an increase in printlet density and printlet weight, hence had a similar impact as compared to laser power density. Moreover, as per Equation 3, the absorptivity of the material is directly proportional to the laser power density, so as per the explanation provided by Fina and colleagues (2018) i.e., a higher amount of Candurin® would lead to a higher energy input that should, in turn, lead to an increase in the hardness of the tablets. However, the contrary was observed where an increase in the amount of Candurin® reduces the hardness of the printlet. This is because unlike photo-absorbing polymers such as polyamides (PA-12) designed for SLS printing in the case of pharmaceutical blends where polymers do not absorb the laser directly, photo absorbing species like Candurin® acts as a conducting excipient, which in-turn causes the thermal transition of the polymer, resulting in sintering. Thereby increasing the amount of Candurin® at the cost of Kollidon® VA64 led to a reduction in the hardness and weight of the printlet. These findings add to the current understanding of the SLS process because properties such as weight critically influence the dose of the printlet, and hardness impacts the stability and performance of the dosage forms.

## 5 Conclusion

In this study, we have demonstrated a straightforward UV-visible spectrophotometric technique which is an efficient pre-formulation tool for predicting the interaction of drugs with the laser source in an SLS process. Further, incorporating a photo-absorbing species such as Candurin® can protect photo-sensitive drugs from degradation, whereas increasing the amount (wt%) of Candurin® at the expense of the polymer reduces the hardness and weight of the printlet. Both laser speed and surface temperature are responsible for the total energy of the surface, where an increase in surface temperature increases the total energy and an increase in laser speed reduces the total energy. High total energy led to an increase in degradation, hardness, printlet weight, and amorphous conversion of the printlet. We demonstrated that an optimum combination of all these factors can be used to process a photo-sensitive drug by SLS. Selective laser sintering has gained much attention in 3D printing pharmaceutical dosage forms, however understanding of the process and its influence on the drug properties are currently lacking. This study demonstrates the importance of performing preformulation testing, and insights on the process and formulation variables on the degradation which can be used to predict, identify and minimize SLS process-induced degradation of a photo-sensitive drug.

## Acknowledgements

The authors would like to acknowledge the Mass Spectrometry Facility at The University of Texas at Austin (https://sites.google.com/site/utaustinmassspec/) and specially Dr. Kristin Blake for access to instrument and resources.

